# Genome expansion by a CRISPR trimmer-integrase

**DOI:** 10.1101/2023.01.23.522648

**Authors:** Joy Y. Wang, Owen T. Tuck, Petr Skopintsev, Katarzyna M. Soczek, Gary Li, Basem Al-Shayeb, Julia Zhou, Jennifer A. Doudna

**Author notes:** These authors contributed equally to this work (Joy Y. Wang, Owen T. Tuck, Petr Skopintsev).

## Abstract

CRISPR-Cas adaptive immune systems capture DNA fragments from invading mobile genetic elements and integrate them into the host genome to provide a template for RNA-guided immunity. CRISPR systems maintain genome integrity and avoid autoimmunity by distinguishing between self and non-self, a process for which the CRISPR-Cas1:Cas2 integrase is necessary but not sufficient. In some microbes, the Cas4 endonuclease assists CRISPR adaptation, but many CRISPR-Cas systems lack Cas4. We show here that an elegant alternative pathway employs an internal exonuclease to select and process DNA for integration using the protospacer adjacent motif (PAM). A natural Cas1:Cas2-exonuclease fusion (trimmer-integrase) catalyzes coordinated DNA capture, trimming and integration. Five cryo-EM structures of the CRISPR trimmer-integrase, visualized both before and during DNA integration, show how asymmetric processing generates size-defined, PAM-containing substrates. Before genome integration, the PAM sequence is released by Cas1 and cleaved by the exonuclease, marking inserted DNA as self and preventing aberrant CRISPR targeting of the host. Together, these data support a model in which CRISPR systems lacking Cas4 use fused or recruited exonucleases for faithful acquisition of new CRISPR immune sequences.

## Introduction

Prokaryotes use CRISPR-Cas (clustered regularly interspaced short palindromic repeats, CRISPR associated) adaptive immune systems to create a sequential genetic record of infection^1^. Transcription and processing of CRISPR sequence arrays consisting of short repeats and ~30 base pair (bp) foreign DNA-derived spacers^2–5^ yield mature CRISPR RNAs (crRNAs) that guide CRISPR-associated (Cas) protein recognition and cutting of complementary genetic material^6–11^. CRISPR-based immunity depends on acquisition of new spacer sequences by the Cas1:Cas2 complex, an integrase responsible for selecting DNA substrates and inserting them into the host CRISPR array^12–14^. The heterohexameric Cas1_4_:Cas2_2_ recognizes DNA fragments (protospacers) consisting of a ~30 bp segment with short 3’ single-stranded DNA (ssDNA) overhangs^14–16^. In many CRISPR systems, efficient protospacer selection requires a flanking 2-5 bp sequence known as the protospacer adjacent motif (PAM) that marks the DNA as foreign. Following DNA capture but before host genome integration, the PAM is removed to prevent aberrant CRISPR-mediated cleavage by Cas9, Cascade or Cas12 and ensure functional crRNA biosynthesis^17–20^. Additionally, the PAM helps CRISPR enzymes rapidly query target sequences^21^ and orchestrates the directionality of spacer insertion^22–24^.

Diverse mechanisms of PAM selection and removal underscore the importance of the PAM for maintaining both adaptive immunity and genome integrity during CRISPR sequence acquisition^23,25–29^. In CRISPR systems including type II-B, some type V, and type I-A, -B, -C, -D, and -G, the Cas4 endonuclease performs PAM selection and processing^26,30–33^. However, ~40%of CRISPR subtypes lack Cas4^25^. In Cas4-less systems such as the type I-E system in the common laboratory *E. coli K12* strain, Cas1 contains a PAM-binding pocket which is believed to play a role in protospacer precursor (prespacer) selection^16,28^. However, whether Cas1 cleaves the PAM in a similar manner to Cas4 or relies on host nucleases to perform this function remains unclear^16,28^. Recent *in vitro* studies identified exonucleases capable of aiding Cas1:Cas2 in prespacer substrate trimming to the correct size for integration^23,28,34^. There are also systems containing a natural Cas2-DEDDh exonuclease fusion^29,35^, implying a functional link between exonucleases and the CRISPR integrase. These systems provide a model for studying the coordination between host exonucleases and the CRISPR integrase during prespacer processing in CRISPR systems that lack Cas4.

Here, we reconstitute CRISPR sequence capture, processing, and integration by a naturally occurring *Megasphaera NM10_related* Cas1:Cas2-DEDDh complex. We find that the Cas1:Cas2-DEDDh complex preserves the PAM during prespacer processing and the first step of integration. The PAM is removed before completing full integration. We show that the DEDDh active site, rather than Cas1^16^, is responsible for both initial 3’ overhang trimming and PAM removal. This mechanism is distinct from that of Cas4, which cleaves the PAM endonucleolytically, suggesting a divergent role for host exonucleases in PAM processing^28^. The integrase regulates DEDDh exonuclease activity by a ruler-guided, ‘gatekeeping’ mechanism that coordinates PAM processing and defines the length of integrated DNA. Cryo-electron microscopy (cryo-EM) structures of Cas1:Cas2-DEDDh bound to prespacer DNA with or without the PAM show how Cas1:Cas2 recognizes the sequence and protects it from DEDDh-mediated DNA trimming. Conformational analysis of half-integration structures suggests that, once anchored into the CRISPR array, DNA bending engages the C-terminal region of Cas1, which in turn exposes the PAM for removal, allowing full integration to occur. Our findings provide a general mechanism for exonuclease-assisted PAM processing and show that CRISPR systems evolved diverse mechanisms of prespacer processing to produce robust immunity against foreign invaders while avoiding autoimmunity.

## Results

### Cas1:Cas2-DEDDh generates CRISPR integration substrates

The adaptive capacity of CRISPR systems relies on the recognition, capture and processing of suitable DNA integration substrates from foreign sources (Fig. 1a). These integration substrates (prespacers) require nucleolytic processing to generate short fragments of uniform length. To investigate the role of the predicted exonuclease domain of Cas2-DEDDh in CRISPR prespacer processing, we expressed and purified Cas1 and Cas2-DEDDh from a type I-E *Megasphaera NM10_related* CRISPR system and tested their ability to process DNA substrates *in vitro* (Fig. 1b). Based on the size of the spacers in the *Megasphaera* CRISPR array and the substrate preferences of the closely related type I-E *E. coli* Cas1:Cas2 integrase, the preferred integration substrate is likely a 23-bp DNA duplex with 5-nt single-stranded 3’ overhangs^15,16^. To test DNA processing activity, we assayed Cas1 and Cas2-DEDDh trimming activity using 5’ fluorophore-labeled prespacer substrates containing a 23-bp duplex region and extended single-stranded 3’ overhangs of varying lengths. Although Cas1 and Cas2-DEDDh each exhibit nuclease activity in isolation, yielding distinct products without apparent functional relevance, only the Cas1:Cas2-DEDDh complex generates substrates equivalent in size to the spacer lengths observed in the host CRISPR array. Processing trims the 3’ overhangs of both strands to produce a 23-bp duplex with 5-6 nt single-stranded 3’ overhangs (Fig. 1b). Repetition of this assay with varying duplex lengths showed that the integrase requires a 23-bp duplex for functional processing (Extended Data Fig. 1a, b). We next tested whether the DEDDh active site is responsible for processing activity by using a catalytically inactive DEDDh mutant, Cas2-DEDDh(D132A), in complex with Cas1. DNA cleavage assays indicated that the mutant complex does not process prespacers (Fig. 1c). Taken together, these data demonstrate that the complete Cas1:Cas2-DEDDh complex is necessary for prespacer processing, with the DEDDh active site providing the relevant nucleolytic activity.

**Figure 1.**
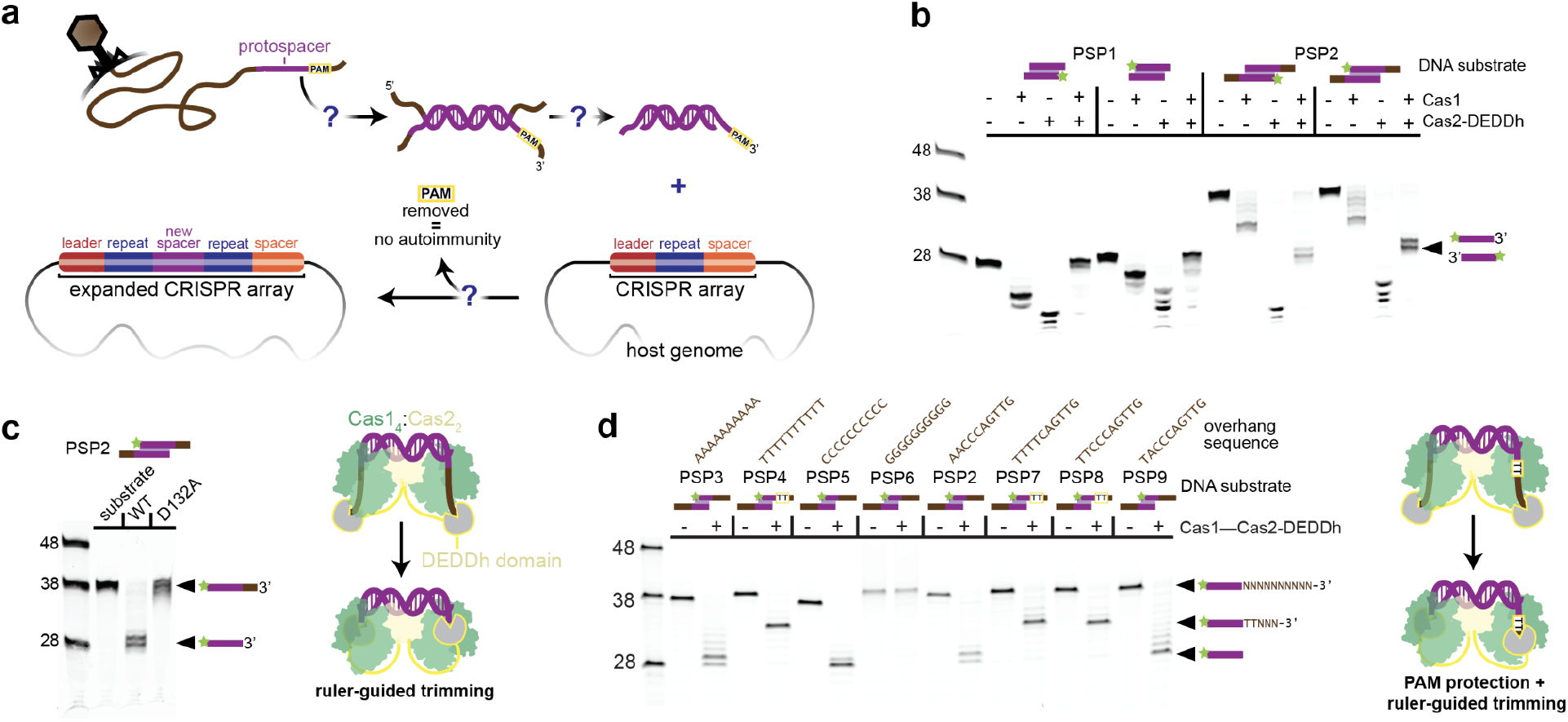
Cas1:Cas2-DEDDh processes prespacers to the correct size for integration and protects a TT PAM. **(a)** Open questions in CRISPR adaptation. **(b)** Processing of fluorescently labeled prespacer substrates with a 23-bp duplex and different overhang lengths by Cas1:Cas2-DEDDh down to 5-6 nt single-stranded 3’ overhangs. **(c)** Wild-type (WT) and mutant Cas1:Cas2-DEDDh (DEDDh-D132A) prespacer processing. **(d)** Processing of substrates with variable 3’ overhangs and model of observed PAM protection and ruler-guided trimming by DEDDh domain.

Time-course assays suggest similar processing efficiencies for prespacer substrates with varying overhang lengths (Extended Data Fig. 2a-d). Fluorescently labeled prespacers were incorporated into an integration target plasmid (pCRISPR) containing a shortened version of the natural *Megasphaera* CRISPR array. Kinetic analysis implies higher relative integration efficiency with the canonical substrate (23-bp duplex with 5-nt single-stranded 3’ overhangs) compared to prespacers with extended overhangs. Reaction with the canonical substrate generated ligation products after 2 min, while prespacers with extended overhangs required 10 min for detection. These data suggest that Cas1:Cas2-DEDDh provides a molecular ruler against which DEDDh trims prespacers to the integration-competent size prior to ligation into the CRISPR array.

To determine the effect of the PAM on prespacer processing, we varied the PAM region, generating a small prespacer library against which the PAM could be inferred (Fig. 1d, Extended Data Fig. 3a,b). We determined that Cas1:Cas2-DEDDh recognizes 5’-TT in the PAM position. In the absence of a TT PAM, DEDDh trims prespacer strands to the integration-competent size (28 nt). However, the presence of a TT PAM in the correct position (nucleotide positions 29 and 30 relative to the 5’ end) results in partial trimming of the PAM-containing strand, precisely 3 nt away from the PAM, yielding a 33 nt product. We hypothesized that partial trimming was the result of sequestration by a PAM binding pocket in Cas1 which protects the PAM and 3 adjacent nucleotides from nuclease activity^16^.

### Cas1 binds the PAM and blocks DEDDh trimming during prespacer processing

We next sought to elucidate the structural basis for prespacer processing and PAM protection. Cryo-EM was used to solve 3.1 Å and 2.9 Å resolution structures of Cas1:Cas2-DEDDh complexed with prespacer substrates with and without a TT PAM, respectively (Fig. 2a-d). In these structures, the *Megasphaera* Cas1:Cas2 retains the canonical heterohexameric architecture^36^, with two Cas1 dimers (denoted Cas1 and Cas1’, a or b subunit) bridged by a central Cas2 dimer (Fig. 2a). A 23-bp duplex sits atop the complex, while 5 nt single stranded 3’ overhangs extend into the clefts formed at opposite Cas1 interfaces. The DEDDh domain could not be resolved in the PAM-absent dataset, presumably due to a high degree of conformational flexibility conferred by the 38 amino acid linker between DEDDh and Cas2, and because all 3’ ends are buried within the complex, protected from exonuclease activity (Fig. 2c). In the PAM-containing dataset, however, the DEDDh domain was resolved by iterative classification and three-dimensional refinement (see Extended Data Fig. 5 and the Methods section).

**Figure 2.**
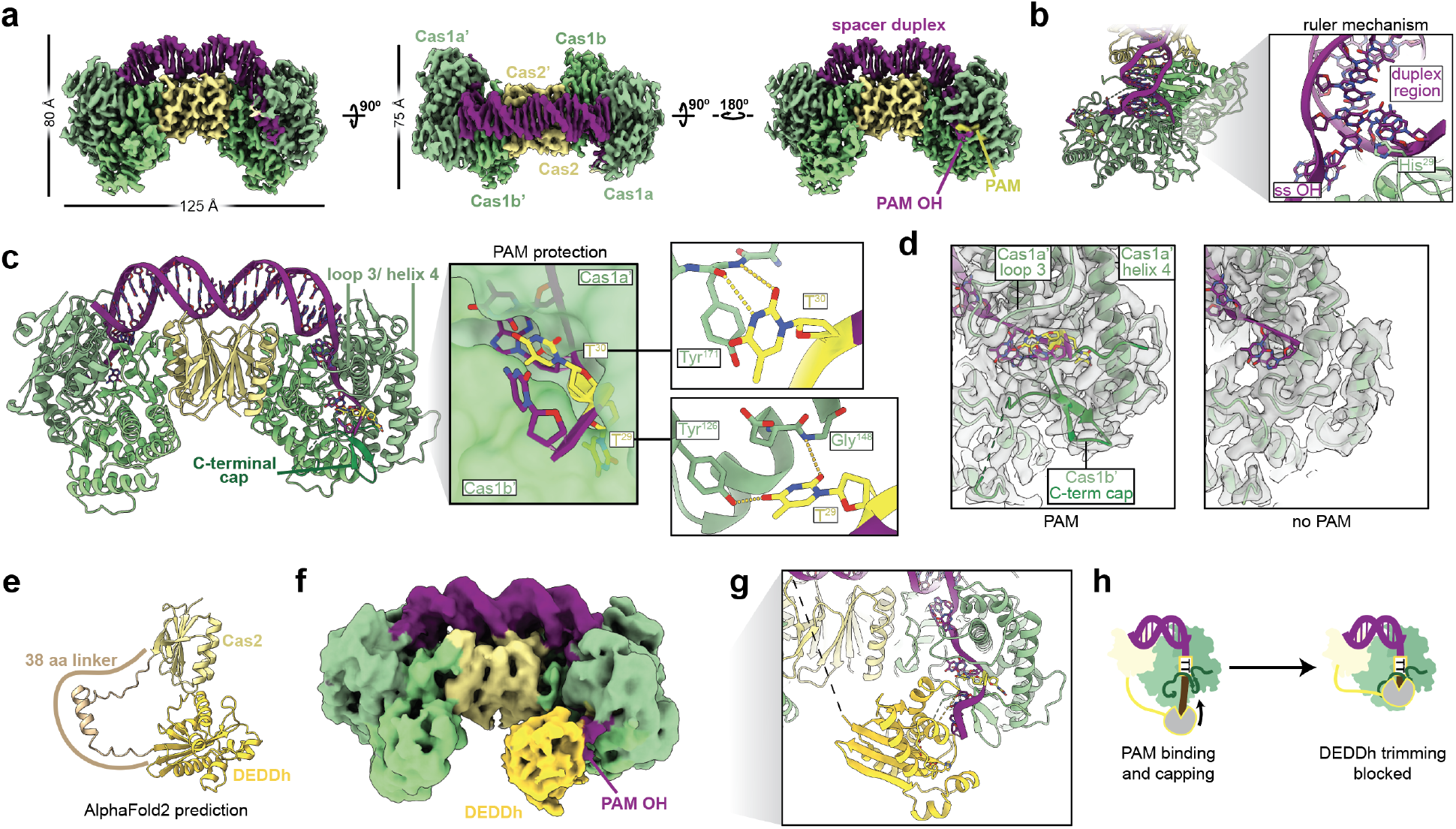
Molecular detail of Cas1:Cas2-DEDDh during prespacer processing. (**a**) Orthogonal views of the final cryo-EM densities for Cas1:Cas2-DEDDh bound to a prespacer containing a phosphorothioated TT PAM (threshold: 0.200). (**b**) Structure of the PAM-bound Cas1:Cas2-DEDDh depicting one of two His^29^ residues dictating duplex length. (**c**) Left, structure viewed from the PAM side. Center, surface depiction of the cleft between Cas1a’ and Cas1b’. Left, sequence specific contacts made with each PAM thymine. (**d**) Comparison of PAM and non-PAM densities with atomic models overlaid at threshold: 0.200. (**e**) AlphaFold 2 prediction of the structure of Cas2-DEDDh. (**f**) Side view of unsharpened cryo-EM density where DEDDh was resolved (threshold: 0.033). (**g**) Hybrid structure containing the DEDDh domain with detail at the PAM-DEDDh interface, with catalytic DEDDh residues shown. Black dashed line represents the unstructured linker between Cas2 and DEDDh domains. (**h**) Model for PAM protection.

Cas1:Cas2-DEDDh dictates prespacer duplex length with an internal ruler that ensures each spacer in the CRISPR array is equivalent in length (Fig. 1b, c). Dual Cas1 His^29^ histidyl residues measure out a 23-bp DNA duplex by π-stacking with the terminal base pairs of the double stranded region, clasping the prespacer strands and marking the start of the single stranded 3’ overhang (Fig. 2b). Biochemical processing assays demonstrate that different duplex lengths are not well tolerated by the complex (Extended Data Fig. 1a, b).

Processing experiments suggest the PAM is protected initially from DEDDh-mediated prespacer processing (Fig. 1b-d). To understand how the integrase sequesters the PAM, we performed cryo-EM analysis on the integrase complex bound to a PAM-containing prespacer DNA molecule with a phosphorothioate backbone modification at the predicted site of DEDDh activity, with the intention of stalling processing. Sequence-specific interactions with loop 3 and helix 4 in the Cas1a’ subunit in the resultant density rationalize PAM recognition (Fig. 2c). The first PAM thymine is buried in a deep pocket in Cas1a’, where hydrogen bonds formed with Tyr^126^ and Gly^148^ may enhance binding affinity (Fig. 2c, right-top). The second PAM thymine π-stacks with Tyr^171^, which positions T^30^ to hydrogen bond with the Tyr^171^ backbone amide nitrogen (Fig. 2c, right-bottom). When the PAM is absent, substrates are fully trimmed, underscoring the necessity of sequence-specific interactions for asymmetric trimming and PAM protection (Fig 1d). Moreover, a beta hairpin in the C-terminal region of Cas1b’ ‘caps’ sequestered nucleotides. Loop 3, helix 4, and the C-terminal cap are absent in the PAM-deficient density, suggesting these structural motifs participate in PAM protection (Fig. 2d).

The DEDDh domain was not visible in the initial PAM-containing structure, raising the question of how the integrase engages the DNA for ruler-guided trimming of sequestered nucleotides. In an effort to resolve the DEDDh density, we iteratively classified and refined the PAM-containing particle stack and found a density corresponding to DEDDh in a small subset of the total particle ensemble (Fig. 2f, Extended Data Fig. 5). Key features include a large protrusion only visible on the PAM side of the complex and an extended density attributable to additional phosphorothioate-containing nucleotides 3’ to the PAM (Fig. 2f). Because the protuberance was low resolution and unsuitable for molecular modeling, the model predicted by AlphaFold 2 for the DEDDh domain was docked into the density (Figs. 2f, g). The resulting hybrid model illustrates dynamics of DEDDh trimming and PAM protection. Catalytic DEDDh residues are poised to exonucleolytically cleave the prespacer overhang, but the PAM binding pocket and the C-terminal loop occlude DEDDh procession, blocking cleavage of the PAM and 2-3 additional nucleotides (Fig. 2h). Despite the high local concentration of nonspecific exonuclease relative to the substrate, this process is precise, in concordance with biochemical evidence (Figs. 1b; 2g). The limited electron density visible for DEDDh in the structurally resolved ensemble is consistent with its intrinsic flexibility. A natural consequence of protection is that the PAM must be cleaved downstream in the process of CRISPR adaptation.

### DEDDh trims the PAM after DNA half-integration

Although there is evidence for PAM protection during prespacer processing, the PAM must be removed prior to insertion into the CRISPR array to avoid autoimmunity. We analyzed Cas1:Cas2-DEDDh processing of DNA substrates designed to mimic intermediates of integration into the CRISPR array to determine the mechanism of PAM removal and resolve dynamics of the complex at the integration target site^28,31^. Two substrates that mimic probable half-integration intermediates - the pre-PAM processing intermediate and the post PAM-processing intermediate (Fig. 3a, half-site substrates 1 and 2, respectively) - were synthesized and assayed in reactions with wildtype and catalytically-inactivated DEDDh complexes. Time-course reactions with half-site substrate 1, which contains the unprocessed PAM, resulted in a 100 nt band corresponding to full-site integration (Fig. 3a). The same band was absent when DEDDh was catalytically inactive. Reactions with PAM-deficient half-site substrate 2 yielded the 100 nt full-site integration product for both the WT and dead DEDDh complexes. Products derived from half-site substrates 1 appear the same size as the full-site integration products generated from half-site substrate 2 (100 nt). These data suggest that, in the wildtype reaction, the PAM is fully removed prior to full-site integration. The lower intensity of the full-site integration product generated from half-site substrate 1 compared to that of half-site substrate 2 may be a result of inefficient PAM removal, also observed in kinetics assays (Extended Data Fig. 2a, b). The absence of the integrated product strand in the catalytically-inactivated DEDDh condition suggests the DEDDh active site executes PAM processing. Notably, PAM processing is necessary for full-site integration, and the integrase generates a precisely defined insertion product size. Hence, nonspecific exonuclease activity generates a ladder of ssDNA overhang fragments in the PAM-containing substrate strand, but only one of these fragment sizes is compatible with full-site integration. This single-nucleotide precision is a result of the Cas1:Cas2 ruler, which simultaneously defines the spacer size and acts as a gate that prevents PAM insertion into the CRISPR array (Fig. 3a). Since Cas1-mediated PAM protection was observed during prespacer processing, it is reasonable to assume that Cas1 releases the PAM for DEDDh trimming while engaged on the CRISPR array.

**Figure 3.**
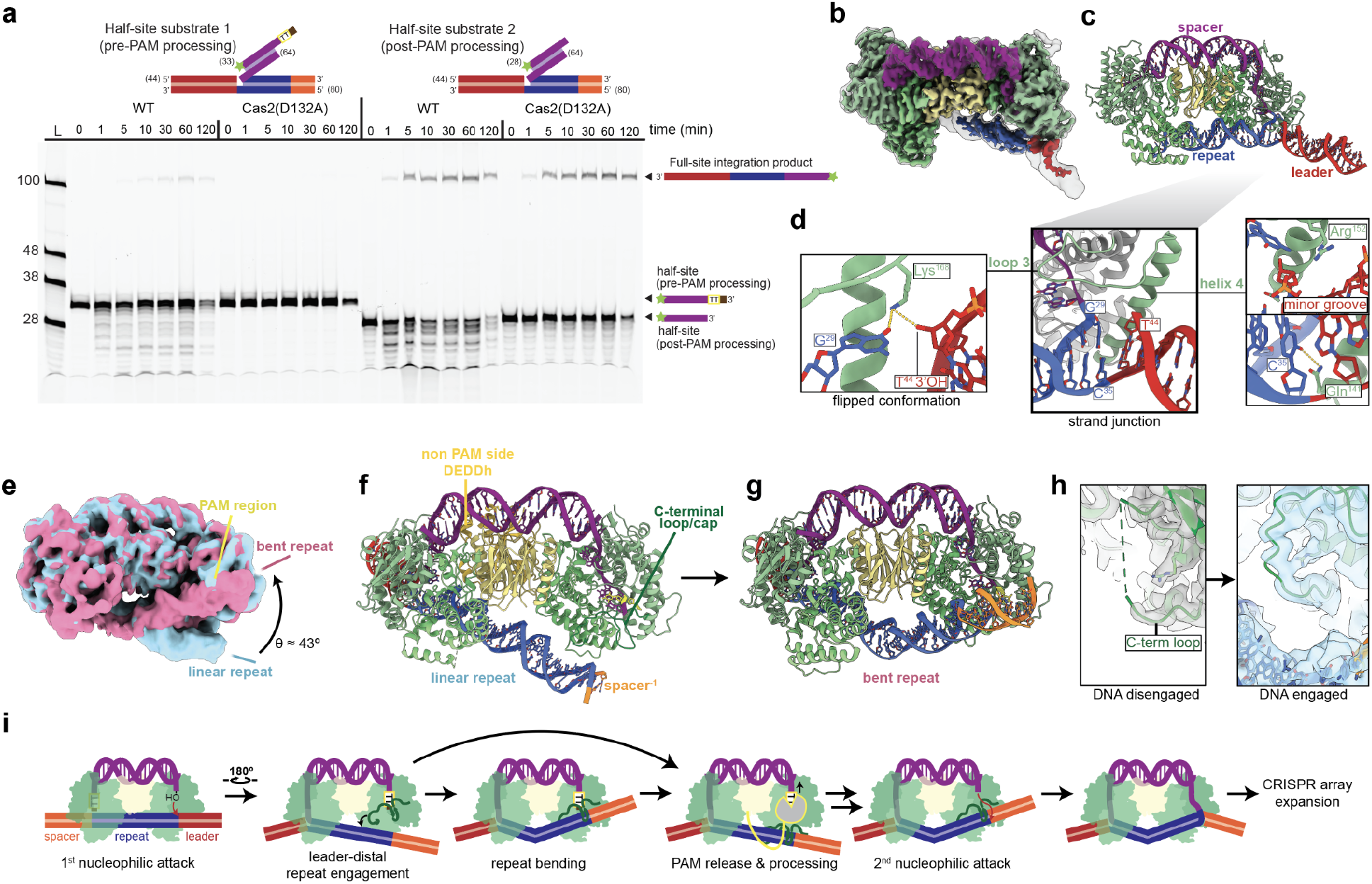
Biochemical and structural analysis of Cas1:Cas2-DEDDh PAM processing. **(a)** Integration reactions by WT and mutant Cas1:Cas2-DEDDh (DEDDh-D132A) half-site integration intermediates. **(b)** Unsharpened (gray transparent, threshold: 0.05) and sharp (color, threshold: 0.19) cryo-EM densities for Cas1:Cas2-DEDDh bound to the PAM-phosphorothioated half-site integration analog. **(c)** Structure of the initial half-site complex. **(d)** Center, detail of the junction of leader, repeat, and spacer. Left, a second ‘flipped’ conformation of G^29^. Right, helix 4 interactions. **(e)** Unsharpened maps of two repeat DNA conformations, linear (blue) and bent (red), both with threshold: 0.05. **(f)** Structure with linear, extended repeat DNA. **(g)** Structure of bent repeat DNA. **(h)** Comparison of C-terminal loop ordering in the prespacer (left) and linear repeat (right) structures. **(i)** Model for PAM gatekeeping facilitated by the C-terminal loop/cap.

Evidence for DEDDh involvement in both prespacer processing and PAM cleavage led us to wonder which molecular cues prompt Cas1 to relinquish the PAM for digestion. Aiming to visualize DEDDh trimming and conformational changes in Cas1, we used cryo-EM to characterize Cas1-Cas2-DEDDh in complex with a DNA half-site analog containing phosphorothioate linkages at the PAM positions (Fig. 3b, Extended Data Fig. 6a-c). Neither the initial 3.1 Å density nor any heterogeneous states detected during cryo-EM data processing had density corresponding to the DEDDh domain on the PAM side of the complex (Fig. 3b). However, DEDDh was observed on the non-PAM side during DNA conformational analysis. We speculate that, in agreement with biochemical data, only the DEDDh domain can trim the PAM, and PAM trimming activity is required for full integration (Fig. 3a). Furthermore, the DEDDh PAM-trimming state may be transient. Phosphorothioate modifications only partially protected the PAM, which may disfavor resolution of the active DEDDh domain at the half-site (Extended Data Fig. 7).

The 3.1 Å half-site structure reveals previously uncharacterized interactions at the first integration strand junction (Fig. 3b-d). After the initial nucleophilic attack, Cas1:Cas2 induces bending of the leader-repeat target DNA^37^. The approximate 26° angle originates at the nick site, positioning repeat DNA between the leader-distal and -proximal Cas1 active sites. Inspection of the strand junction (Fig. 3d, middle) reveals previously undescribed sequence-specific interactions with the first base pair of the CRISPR repeat. Two continuous secondary structures, loop 3 and helix 4 of Cas1a’, dictate these interactions. The spacer-ligated first CRISPR repeat guanine G^29^ was found in two approximately equivalent conformations. In the first conformation, G^29^ forms a canonical base pair with C^35^ (Fig. 3d, middle). In the second conformation, G^29^ ‘flips’ upward, making a specific contact with loop 3 lysine Lys^168^. The lysine also contacts the 3’ hydroxyl of the leader (Fig. 3d, left). Gln^141^, which sits at the base of helix 4, also makes a nucleobase-specific contact with C^35^, the first bottom repeat nucleotide. Helix 4 is well positioned to insert into the minor groove of the leader DNA, but no nucleobase-specific contacts were obvious (Fig. 3d, right). Specific contacts with CRISPR array nucleotides likely play a functional role in insertion targeting.

The 3.1 Å half-site complex reconstruction contains density corresponding only to the leader-proximal region of the CRISPR repeat (Fig. 3b, c). To probe dynamics at the leader-distal region, where PAM processing and subsequent full integration occur, we performed three-dimensional variability analysis (3DVA) on the particle set (Extended Data Fig. 6a, d, e)^38^. 3DVA revealed heterogeneity in the location of the repeat/spacer end, with the CRISPR repeat DNA oscillating between linear and bent conformations (Fig. 3e, Supplementary Video 1). The DEDDh domain was visible only in the linear conformation. Isolation and refinement of particle clusters representing maxima of the reaction coordinate gave linear and bent reconstructions at 4.1 Å and 3.9 Å resolutions, respectively (Fig. 3e-g).

In the linear structure, a Cas1b’ C-terminal loop (L^279^ to S^293^) rich in charged residues is positioned near the major groove adjacent to the second integration target site (Fig. 3f). The corresponding density is absent in the 3.1 Å half-site and 2.9 Å prespacer-bound structures, indicating this loop participates in engagement with the CRISPR repeat on the PAM side (Fig. 2c; 3g, h). A C-terminal cap, which follows the C-terminal loop, protects the PAM and adjacent nucleotides from trimming by DEDDh (Fig. 2h). Intriguingly, the DEDDh domain was visible in the linear structure, but on the non-PAM side of the complex, where no overhang trimming occurs (Fig. 3f). The exonuclease sits in a cavity formed by the interface of the CRISPR repeat DNA, Cas1a/b, and Cas2, where it appears to contact the repeat DNA backbone and the N-terminus of the Cas1b (Fig. 2f, Extended Data Fig. 8). These interactions may stabilize leader-proximal DNA and beneficially constrain bending of the second integration target site towards the Cas1a’ active site while DEDDh is not engaged in trimming.

The bent structure features a pronounced ~43° kink in the center of the repeat region and at the bottom of the holoenzyme (Fig. 4f). Disruption of a single A-T base pair and DNA unwinding appear to accommodate the strain induced by this pitch, though this assignment was made with low confidence due to the low local resolution at the bending site. Bending in the center of the CRISPR repeat symmetrizes the integration complex and draws the repeat/spacer junction towards the Cas1a’ active site. Although PAM nucleotides are still present in this density, likely due to their cleavage-blocking phosphorothioate modifications, the overhang nucleotides typically sequestered during ruler guided trimming are absent, suggesting that 3’ trimming occurs in an intermediate step between linear and bent states. Only the first three nucleotides under the C-terminal loop of Cas1b’ could be assigned, and the C-terminal cap density was largely unstructured. These observations, combined with the proximity of the second integration target site in the bent structure, suggest that engagement of the CRISPR DNA by the C-terminal loop induces uncapping of previously sequestered PAM nucleotides, followed by DEDDh- or host exonuclease-mediated trimming of the exposed 3’ end to the ruler-defined length. Once the PAM is trimmed, full integration occurs (Fig. 3a). Structural and biochemical analyses imply a general mechanism of sequential PAM protection and cleavage, or PAM gatekeeping, which ensures that the PAM-deficient protospacers integrated into the CRISPR array are marked as “self” and are equal in size (Fig. 3i).

**Figure 4.**
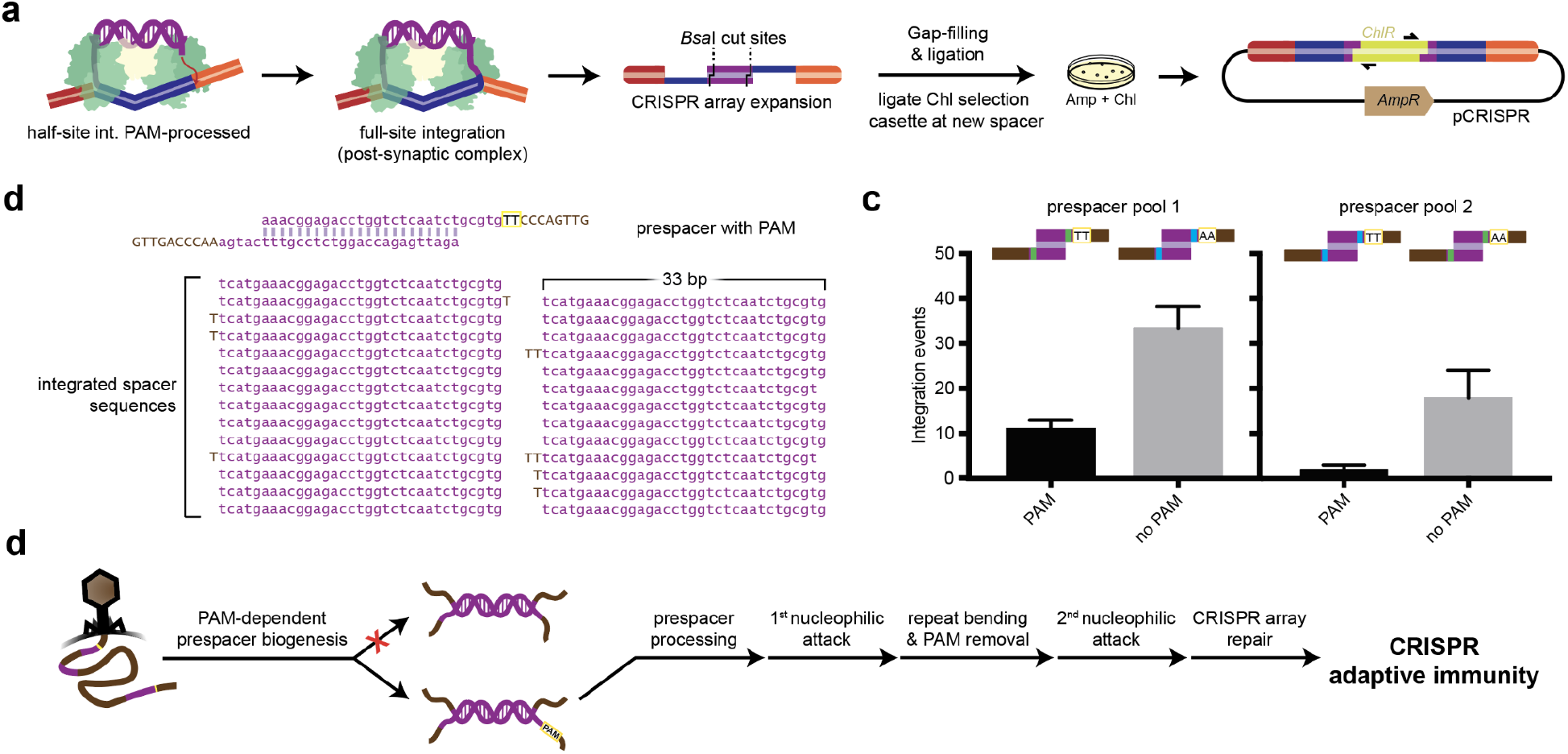
*In vitro* reconstitution of Cas1:Cas2-DEDDh-mediated full-site integration. **(a)** Schematic *in vitro* reconstitution of full-site integration. **(b)** Fully integrated spacer sequences from a prespacer containing the PAM. **(c)** Number of integration events arising from an equimolar pool with or without the PAM. Prespacers are distinguished by internal barcodes which were swapped to remove sequence bias. The mean and standard deviation of three independent replicates are shown. **(d)** General timeline of CRISPR adaptation.

### Reconstitution of full CRISPR array integration

To determine how PAM sequence recognition and gated removal ensures accurate DNA integration, we reconstituted CRISPR substrate integration *in vitro*. An unprocessed prespacer (23-bp duplex with 15-nt single-stranded overhangs) containing the TT PAM was combined with Cas1:Cas2-DEDDh and pCRISPR, which contains a shortened version of the *Megasphaera* CRISPR array. Prespacer substrates encoded BsaI restriction sites in the duplex region to enable insertion of a chloramphenicol resistance gene. Upon transformation, only pCRISPR with full prespacer integration confers survival in a double selection assay (Fig. 4a)^39^. Fully integrated sequences provide additional evidence that Cas1:Cas2-DEDDh completes both prespacer processing and integration into the CRISPR array (Fig. 4b). The complex is specific for the CRISPR array and all integration events occur at repeat borders (Extended Data Fig. 9b). However, integration occurs at all three repeats present in the array, without specificity for the leader-proximal repeat, as seen in many CRISPR systems *in vivo^2,40^*. This is consistent with structural data, which only shows specificity for the first base pair of the repeat (Fig. 3d). Excess 3’ overhangs are trimmed to within 1-2 nt of the expected length and the PAM is absent in all integrated sequences (Fig. 4b), in agreement with evidence at the level of the half-site (Fig. 3a, i). While the Cas1:Cas2-DEDDh complex initially recognizes and protects the PAM from trimming, it removes the PAM prior to full integration. Reconstitution experiments provide complementary evidence for an alternative mechanism for PAM processing compared to Cas4, which uses a sequence-specific mechanism to cleave the PAM endonucleolytically^30,31^.

Although it was hypothesized that delayed PAM trimming aids the complex in orienting the prespacer for integration^28,31^, no orientation bias was observed *in vitro* (Extended Data Fig. 9c). While Cas1:Cas2-DEDDh alone is able to distinguish between the PAM and non-PAM sides of the prespacer (Figs. 1d; 2c), it appears that the complex alone cannot discern the leader-side and spacer-side of the repeat, consistent with cryo-EM results of the half-site intermediate, which show no sequence specificity for the leader (Fig. 3d). We suspect the complex requires additional host factors to correctly orient the spacers *in vivo*. In *E. coli*, integration host factor (IHF) directs the first nucleophilic attack to the leader-side of the repeat through specific contacts with the leader sequence^37,41^. Superimposition of the half-site structure and a *Megasphaera* IHF ortholog onto a structure of the complete IHF-containing integration holo complex further implicates the participation of a host factor in directing CRISPR integration (Extended Data Fig. 10). *In vivo*, the system may have higher specificity for the leading integration target site and utilize delayed PAM processing as the basis for determining the orientation of integration, as is the case in other CRISPR systems^23,28,31^.

To assess the effect of the PAM on integration efficiency, an equimolar mixture of PAM-deficient and PAM-containing prespacers, each containing a pair of identifying internal barcodes, was tested for full-site integration. Surprisingly, out of 95 sequenced colonies, we observed significant enrichment (~3-fold) of integration events from the PAM-deficient prespacer (Fig. 4c). To account for biases resulting from the internal barcode sequences, we generated a second prespacer pool, where barcode pairs were swapped. The second pool also exhibited a significant preference for the PAM-deficient prespacer (Fig. 4c). Lower integration efficiency from the PAM-containing prespacer *in vitro* may stem from additional steps required for PAM removal (Fig. 3i). Moreover, PAM removal is observed following full-site integration, and all spacer sequences are selected according to PAM presence (Fig. 4b). The reduced apparent efficiency of PAM-containing prespacer insertion *in vitro* therefore suggests PAM recognition *in vivo* occurs upstream, during the biogenesis of substrates bound for CRISPR adaptation (Fig. 4d).

Integration reconstitution experiments with pooled prespacers suggest that the integrase may select substrates prior to prespacer processing, ensuring PAM presence (Fig. 4c). We were interested in whether the integrase demonstrates similar stringency for another substrate feature, the canonical prespacer duplex. Accordingly, we tested Cas1:Cas2-DEDDh processing after stepwise addition of the PAM-complementary strand. After incubation of Cas1:Cas2-DEDDh with a single-stranded PAM-containing strand, the labeled, PAM-deficient strand was added. In all reactions, even for the prespacer strands with the tolerated 23-nt complementarity region (Fig. 1b), nonspecific processing of the labeled strand occurs (Extended Data Fig. 10d-f). Thus, Cas1:Cas2-DEDDh likely performs ruler-guided trimming when the 23-bp prespacer is pre-duplexed and not after delayed addition or search for the complementary strand. The strict requirement for substrate size, strandedness, and PAM-presence has implications for open questions in CRISPR substrate biogenesis.

## Discussion

Efficient CRISPR adaptive immunity requires coordination between the CRISPR integrase and host nucleases^23,28^. In this study, we describe mechanisms of prespacer processing and integration in a naturally occurring Cas1:Cas2-DEDDh complex. The trimmer-integrase uses an alternative PAM-processing mechanism compared to the well-studied Cas4 endonuclease. Prior to this work, it was unclear how systems lacking Cas4 process integration substrates. Our data suggest that one evolutionary solution to the problem of selecting, protecting, and then removing the PAM is to use Cas1 rather than an accessory protein for initial PAM protection. Sequestration of defined prespacer sizes via substrate gatekeeping ensures that the PAM is present and that its cognate spacer is functional (Fig. 1d). Once the PAM-containing prespacer is anchored to the host CRISPR array, the PAM is exposed by the Cas1 ‘gate’ and is promptly removed at single nucleotide resolution. We provide a mechanism explaining which structural cues lead to PAM uncapping and removal (Fig. 3i). Binding and bending of leader-distal repeat DNA may lead to disengagement of the C-terminal cap which covers and protects nucleotides. Once exposed, DEDDh completely digests the PAM, generating substrates of the correct size and positions them for nucleophilic attack at the repeat/spacer junction. This sequence ensures that the PAM side integrates second. Bending and unwinding may additionally aid in melting of the repeat strand, which is required for resolution of the post-synaptic complex and concomitant repeat duplication^42^. While the high effective local concentration of exonuclease with respect to the bound prespacer conferred by the Cas2-DEDDh fusion in this system likely improves efficiency, we imagine that host exonucleases can function similarly in *trans*. Cas1:Cas2-DEDDh serves as a general model which describes the role of accessory exonucleases, including those which aren’t fused to the nuclease, in the diverse CRISPR systems lacking Cas4. Our findings advance our understanding of how prespacers are processed and selected for spacer acquisition. We anticipate that our results will be applicable to CRISPR-based technologies which seek to repurpose Cas1:Cas2 for the molecular recording of cellular events, an application challenged both by reliance on host factors such as exonucleases and by uncertainty in prespacer selection^22,43,44^.

While this report represents progress in the understanding of PAM processing immediately before integration, upstream PAM selection and prespacer biogenesis remain mysterious. Unexpectedly, experimental data suggest lower integration efficiency from PAM-containing prespacers and a preference for preformation of the canonical duplex^15^. Together, these data weaken the ‘complement search’ model for CRISPR integration substrate biogenesis, which suggests single stranded DNA derived from foreign sources are captured independently by the integrase complex. Alternatively, Cas1:Cas2 may recognize PAM-containing prespacer-like motifs as DNA reanneals behind repair complexes implicated in CRISPR adaptation such as RecBCD and AddAB, as was recently suggested^33^. The precise mechanistic details of this proposal are unclear. Future experiments might utilize the compact trimmer-integrase presented here to probe open questions in CRISPR substrate biogenesis and achieve total *in vitro* reconstitution of naive CRISPR adaptation.

## Materials and Methods

### Plasmid construction and DNA substrate preparation

To make the target integration plasmid pCRISPR, the leader and the first 3 repeats and spacers of the CRISPR array were ordered as two DNA fragments, which were amplified by PCR and inserted into a pUC19 backbone by Gibson Assembly. DNA oligos used in this study were ordered from Integrated DNA Technologies (IDT). Prespacers and the half-site substrates were formed by heating at 95 °C for 5 min and slow cooling to room temperature in HEPES hybridization buffer (20 mM HEPES, pH 7.5, 25 mM KCl, and 10 mM MgCl_2_). For the half-site substrate, hybridization was carried out with a 1.5-fold excess of the two shortest strands and 1.25-fold excess of the second-largest strand and purified on a 8% native PAGE gel. Sequences of cloning primers and DNA substrates are shown in Table S2.

### Cloning, expression, and purification

The *Megasphaera NM10_related* Cas1 and Cas2-DEDDh genes were codon optimized for *E. coli* expression, ordered as G-blocks, PCR amplified, and cloned separately into a pET-based expression vector with an N-terminal 10xHis-MBP-TEV tag. After transformation into chemically competent Rosetta cells, cells were grown to an OD600 of ~0.6 and induced overnight at 16 °C with 0.5 mM isopropyl-β-D-thiogalactopyranoside (IPTG). Cells were harvested and resuspended in lysis buffer (20 mM HEPES, pH 7.5, 500 mM NaCl, 10 mM imidazole, 0.1% Triton X-100, 1 mM Tris (2-carboxyethyl)phosphine (TCEP), Complete EDTA-free protease inhibitor (Roche), 0.5 mM phenylmethylsulfonyl fluoride (PMSF), and 10% glycerol). After lysis by sonication and clarification of lysate by centrifugation, the supernatant was incubated with Ni-NTA resin (Qiagen). The resin was washed with wash buffer (20 mM HEPES, pH 7.5, 500 mM NaCl, 10 mM imidazole, 1 mM TCEP, and 5% glycerol) and protein was eluted with wash buffer supplemented with 300 mM imidazole. After overnight digestion with TEV protease, the salt concentration was diluted to 300 mM NaCl using ion-exchange buffer A (20 mM HEPES, pH 7.5, 1 mM TCEP, and 5% glycerol) and run through a tandem MBPTrap column (GE Healthcare) and HiTrap heparin HP column (GE Healthcare) to remove the MBP and bind the protein onto the heparin column. The protein was eluted with a gradient from 300 mM to 1 M KCl, concentrated, and purified on a Superdex 200 (16/60) column with storage buffer (20 mM HEPES, pH 7.5, 500 mM KCl, 1 mM TCEP, and 5% glycerol). The same purification protocol was used for Cas1 and Cas2-DEDDh (WT and D132A mutant). Sequences of proteins are shown in Table S1.

### Processing assays

Processing assays were conducted in integration buffer (20 mM HEPES, pH 7.5, 125 mM KCl, 10 mM MgCl_2_, 1 mM DTT, 0.01% Nonidet P-40, and 10% DMSO). Cas1 (4 μM) and Cas2-DEDDh (2 μM) were pre-complexed for 30 min. at 4 °C before addition of fluorescent DNA substrate (312.5 nM) and reacting for 2 hours at 37 °C. Reaction was quenched by addition of 2 vol. quench buffer (95% formamide, 30 mM EDTA, 0.2% SDS, and 400 μg/mL heparin) and heating at 95 °C for 4 min., before analysis on a 14% urea-PAGE gel. Reactions were visualized by Typhoon FLA gel imaging scanner and quantification of intensities was performed with ImageQuantTL 8.2. The percent processing activity is quantified as the ratio of the final product bands’ intensity to the total intensity of all bands in the lane.

### Cryo-EM data acquisition

Cas1-Cas2-DEDDh DNA complexes were formed by mixing 50 μM Cas1, 50 μM Cas2-DEDDh, and 12.5 μM prespacer or half-site DNA and dialyzing for 2 h using a Slide-A-Lyzer MINI Dialysis Device at room temperature. The complex was concentrated to varying concentrations of Cas1:Cas2-DEDDh (see Extended Data Table 3) and purified over a Superose 6 Increase 10/300 GL column. Samples were frozen using FEI Vitrobot Mark IV, cooled to 8°C at 100% humidity. Depending on the sample (see Supplementary Table 3), either carbon 2/2 300 mesh C-flat grids (Electron Microscopy Sciences CF-223C-50) or 1.2/1.3 300 mesh UltrAuFoil gold grids (Electron Microscopy Sciences #Q350AR13A) were glow discharged at 15 mA for 25 s using PELCO easyGLOW. In all cases, a total volume of 4 μL sample was applied to the grid and immediately blotted for 5 s with a blot force of 8 units. Micrographs were collected on a Talos Arctica operated at 200 kV and x36,000 magnification (1.115 A pixel size), in the super-resolution setting of K3 Direct Electron Detector. Cryo-EM data was collected using SerialEM v.3.8.7 software. Images were obtained in a series of exposures generated by the microscope stage and beam shifts.

### Cryo-EM data processing

All datasets were collected with varied tilt angle, amount of movies, and defocus range (see Supplementary Table 3 and Extended Data Figures 4, 5, 6). Data processing was performed in cryoSPARC v3.2.0, v3.3.1, and v4.1.1^45^. Movies were corrected for beam-induced motion using patch motion correction, and contrast transfer function (CTF) parameters were calculated using patch CTF.

PAM-deficient prespacer bound Cas1:Cas2-DEDDh map was obtained by an iterative process. In the first round 569 particles were picked manually from 37 micrographs and submitted for Topaz training^46^. The resulting Topaz model was used to pick particles from the micrographs, and a total of 460,631 particles were extracted with bin factor 2, and applied to 2D classification. Following the selection of the best classes, 410,757 particles were used for *ab initio* reconstruction and subsequent heterogenous refinement, with three classes. All particles were used for non-uniform (NU) map refinement^47^, and an initial complex map was obtained. Following 2D classification of particles from the initial NU refinement model 38,342 particles from the classes with isotropic orientations were selected and subject to the second round of Topaz training. A new Topaz model was used with a total of 956 curated micrographs, and the entire process was repeated twice with particles from the best heterogeneous refinement class for subsequent NU refinement and Topaz training. The final map with the best electron density for the PAM-deficient prespacer bound Cas1:Cas2-DEDDh complex was obtained from 461,266 particles and was refined with non-uniform refinement to 3.1 Å.

For the PAM-containing prespacer bound Cas1:Cas2-DEDDh, a single round of Topaz training was applied. Following the initial exposures curation which yielded 591 best quality micrographs, 6302 particles were manually picked and subject to Topaz training job. The Topaz model was applied to an expanded set of 1184 curated micrographs, and resulted in extraction of 3,101,776 particles. After *ab initio* reconstruction and heterogenous refinement of the particles, with three classes, the 1,420,721 particles set constituting the best class were processed with non-uniform refinement. As a result, a 2.9 Å density for PAM-containing prespacer bound Cas1-Cas2 complex was obtained.

For resolving the DEDDh density in the latter dataset, the *Ab initio* class particles used for the latter density reconstruction, 1,331,357 in total, were were applied to a 2D classification job, and 228,220 particles were selected in classes with apparent DEDDh density. Following *ab initio* refinement with three classes, particles from the best class were subject to another round of 2D classification, and 109,912 particles with more pronounced DEDDh density were selected, and re-extraction with 320 pixel box size (in all other cases, 480 pixel boxes were used for the extraction jobs). As a result of the final 2D classification round, 49,560 particles with the best DEDDh density were selected, re-extracted with standard box dimensions, and subject to *ab initio* refinement, with one class, and non-uniform refinement. As a result, a 3.5 Å complex map with the DEDDh exonuclease density was obtained, with a total of 49,383 particles used for reconstruction.

For half-site DNA-bound Cas1:Cas2-DEDDh, the Topaz model from the PAM-containing prespacer was applied to 2,810 micrographs selected after manual curation. The 2,448,888 resultant particles were subdivided using 2D classification, and the 25 best classes were selected, resulting in 1,836,610 particles. These particles were subjected to *ab initio* reconstruction with three classes. The best class containing 1,048,353 particles was refined using non-uniform refinement to yield to the 3.1 Å half-site map.

To observe DNA dynamics in the Cas1:Cas2-DEDDh half-integration complex, we performed 3-D variability analysis (3DVA)^38^ on a subset of particles selected and refined from 2-D classification with DNA visible on the leader-distal side of the complex (1,048,353 particles). The filter resolution was 6 Å and the number of modes was 3. To generate Supplementary Video 1, the 3DVA output mode was set to simple and 20 frames, then UCSF ChimeraX was used to generate a “vseries.” Next, the 3DVA output mode was set to cluster and the number of clusters to 20. Each resulting cluster was individually inspected and two clusters representing maxima of DNA motion along the pitch axis were chosen. The linear structure was derived from 32,722 particles and was subjected to non-uniform refinement to give the final 4.1 Å map. The bent structure resulting from initial 3DVA clustering was improved by repetition of the 3DVA workflow with the complete particle set obtained by Topaz picking, then selection and non-uniform refinement of the cluster representing leader-distal DNA in the most bent conformation (53,545 particles total), yielding the final 3.9 Å map.

### Model building and refinement

The initial models of the Cas1 and Cas2-DEDDh were obtained with the AlphaFold 2 program^48^. To build the model of Cas1:Cas2-DEDDh bound to a prespacer with TT PAM complex, the predicted Cas1, Cas2 monomers were docked independently into the corresponding map with the fitmap tool in UCSF ChimeraX v1.2.5^49^. The DNA models were built de novo. The complex model was refined using rounds of real-space refinement and rigid body fit tools in Coot v0.9.4.1^50^, and real_space_refine tool in Phenix v1.19.2-4158^51^, using secondary structure, Ramachandran, and rotamer restraints. This complex model served as an initial model for other Cas1:Cas2 structures, which were refined in an analogous manner.

### Ligation assays with pCRISPR integration target plasmid

Ligation assays were conducted in integration buffer (20 mM HEPES, pH 7.5, 125 mM KCl, 10 mM MgCl_2_, 1 mM DTT, 0.01% Nonidet P-40, and 10% DMSO). Cas1 (4 μM) and Cas2-DEDDh (2 μM) were pre-complexed for 30 min. at 4 °C before addition of DNA substrate (312.5 nM) and integration target pCRISPR (20 ng/mL, ~10 nM) and reacting for 2 hours at 37 °C. Reaction was quenched with 0.4% SDS and 25 mM EDTA, treated with proteinase K for 15 min at room temperature, and then treated with 3.4% SDS. Reactions were analyzed on a 1.5% agarose gel and visualized by Typhoon FLA gel imaging scanner.

### Full-site integration assays

Integration assays (50 uL reactions) were conducted in integration buffer (20 mM HEPES, pH 7.5, 125 mM KCl, 10 mM MgCl_2_, 1 mM DTT, 0.01% Nonidet P-40, and 10% DMSO). Cas1 (4 μM) and Cas2-DEDDh (2 μM) were pre-complexed for 30 min. at 4 °C before addition of DNA substrate containing BsaI cut sites (312.5 nM) and reacting for 15 minutes, followed by the addition of the integration target pCRISPR (20 ng/mL, ~10 nM) and incubating for 2 hours at 37 °C. Products were purified with a DNA Clean and Concentrator 5 kit (Zymo Research) and eluted with 6 μl water. A gap-filling reaction (20 μl total, 37 C for 30 min) was conducted with the purified integration products as described by a previous study^39^: 6 μl purified acquisition reaction, 6.5 μl water, 2 μl 10x Taq DNA Ligase buffer (NEB), 2 uL dNTP Solution Mix (10 mM stock, NEB), 2 μl Taq DNA ligase (80 U, NEB), and 1 μl T4 DNA Polymerase (1 U, NEB). Gap-filling reactions purified using the Zymo Research kit and eluted with 6 uL water. A Golden Gate-compatible chloramphenicol selection cassette was generated by PCR with primers encoding BasI cut sites and purified using the Qiagen MinElute PCR Purification Kit. The sequences of primers used are shown in Table 4.2. A Golden Gate cloning reaction was performed with the purified, gap-filled integration products and chloramphenicol selection cassette following a standard BsaI assembly protocol. The products were purified using the Zymo Research kit and eluted with 6 uL water and 1 uL was electroporated into DH10B cells (NEB). Electroporated cells were recovered in 975 uL of LB and plated on LB agar containing carbenicillin (100 μg/ml) and chloramphenicol (25 μg/ml). Of the surviving colonies, 95 were sequenced using Sanger Sequencing and sequences were analyzed using SnapGene Version 5.0.8.

### CRISPR locus bioinformatic analysis

Cas2-DEDDh-containing loci from metagenomic data were identified by determining genomes that contained a CRISPR locus via CRISPRDetect, and coding sequences within 5kb of the array were extracted^52^. A DEDDh HMM model was built from BLAST searches against the NCBI nr database that were manually verified^53^. The coding sequences were searched against the DEDDh model using hmmsearch with E < 1 × 10-5^53^. Matches that also contained credible hits to Cas1 and neighboring other Cas proteins were shortlisted for this work. A preliminary Cas2-DEDDh model was computed using AlphaFold 2 to aid in structure-building^48^.

### Statistics and reproducibility

For biochemical experiments, results represent gels of the highest quality. All experiments were generally performed at least in duplicate, though not in the exact same format. For full-site experiments, we typically performed assays in biological triplicate to ensure reproducibility. All individual data points are displayed on figure panels.

## Acknowledgments

The authors thank members of the Doudna lab for insights and N. Prywes for critical reading of this manuscript. Financial support was provided by grants from the National Science Foundation. J.Y.W. and G.L. were supported by U.S. National Science Foundation Graduate Research Fellowships (DGE 1752814 and DGE 2146752). P.S. was supported by the Swiss National Science Foundation Mobility fellowship (P2EZP3_195621). J.A.D. is a Howard Hughes Medical Institute investigator.

## Author Contributions

J.Y.W. conceived of this project and designed experiments with O.T.T., P.S., G.L., and J.A.D.. J.Y.W., O.T.T., G.L., and J.Z. performed protein purification and *in vitro* biochemistry. P.S. performed cryo-EM data collection with K.M.S. and O.T.T.. P.S., O.T.T., and K.M.S. generated cryo-EM maps and built atomic models with input from J.Y.W.. B.Al.-S. identified the CRISPR-Cas locus of interest and performed bioinformatics experiments. O.T.T., J.Y.W., and J.A.D. wrote the manuscript with input from all authors.

## Additional Information

Supplementary Information is available for this paper.

## References

1. Labrie, S. J., Samson, J. E. & Moineau, S. Bacteriophage resistance mechanisms. Nat. Rev. Microbiol. 8, 317–327 (2010).

2. Barrangou, R. et al. CRISPR provides acquired resistance against viruses in prokaryotes. Science 315, 1709–1712 (2007).

3. Yosef, I., Goren, M. G. & Qimron, U. Proteins and DNA elements essential for the CRISPR adaptation process in Escherichia coli. Nucleic Acids Res. 40, 5569–5576 (2012).

4. Sternberg, S. H., Richter, H., Charpentier, E. & Qimron, U. Adaptation in CRISPR-Cas systems. Molecular Cell 61, 797–808 (2016).

5. McGinn, J. & Marraffini, L. A. Molecular mechanisms of CRISPR–Cas spacer acquisition. Nat. Rev. Microbiol. 17, 7–12 (2019).

6. Pourcel, C., Salvignol, G. & Vergnaud, G. CRISPR elements in Yersinia pestis acquire new repeats by preferential uptake of bacteriophage DNA, and provide additional tools for evolutionary studies. Microbiology 151, 653–663 (2005).

7. Mojica, F. J. M., Díez-Villaseñor, C., García-Martínez, J. & Soria, E. Intervening sequences of regularly spaced prokaryotic repeats derive from foreign genetic elements. Journal of Molecular Evolution 60, 174–182 (2005).

8. Bolotin, A., Quinquis, B., Sorokin, A. & Ehrlich, S. D. Clustered regularly interspaced short palindrome repeats (CRISPRs) have spacers of extrachromosomal origin. Microbiology 151, 2551–2561 (2005).

9. Brouns, S. J. J. et al. Small CRISPR RNAs guide antiviral defense in prokaryotes. Science 321, 960–964 (2008).

10. Garneau, J. E. et al. The CRISPR/Cas bacterial immune system cleaves bacteriophage and plasmid DNA. Nature 468, 67–71 (2010).

11. Hale, C. R. et al. RNA-guided RNA cleavage by a CRISPR RNA-Cas protein complex. Cell 139, 945–956 (2009).

12. Nuñez, J. K., Lee, A. S. Y., Engelman, A. & Doudna, J. A. Integrase-mediated spacer acquisition during CRISPR–Cas adaptive immunity. Nature 519, 193–198 (2015).

13. Wright, A. V. & Doudna, J. A. Protecting genome integrity during CRISPR immune adaptation. Nat. Struct. Mol. Biol. 23, 876–883 (2016).

14. Xiao, Y., Ng, S., Nam, K. H. & Ke, A. How type II CRISPR–Cas establish immunity through Cas1–Cas2-mediated spacer integration. Nature 550, 137–141 (2017).

15. Nuñez, J. K., Harrington, L. B., Kranzusch, P. J., Engelman, A. N. & Doudna, J. A. Foreign DNA capture during CRISPR–Cas adaptive immunity. Nature 527, 535–538 (2015).

16. Wang, J. et al. Structural and mechanistic basis of PAM-dependent spacer acquisition in CRISPR-Cas systems. Cell 163, 840–853 (2015).

17. Deveau, H. et al. Phage response to CRISPR-encoded resistance in Streptococcus thermophilus. J. Bacteriol. 190, 1390–1400 (2008).

18. Mojica, F. J. M., Díez-Villaseñor, C., García-Martínez, J. & Almendros, C. Short motif sequences determine the targets of the prokaryotic CRISPR defence system. Microbiology 155, 733–740 (2009).

19. Gleditzsch, D. et al. PAM identification by CRISPR-Cas effector complexes: diversified mechanisms and structures. RNA Biol. 16, 504–517 (2019).

20. Marraffini, L. A. & Sontheimer, E. J. Self versus non-self discrimination during CRISPR RNA-directed immunity. Nature 463, 568–571 (2010).

21. Cofsky, J. C., Soczek, K. M., Knott, G. J., Nogales, E. & Doudna, J. A. CRISPR-Cas9 bends and twists DNA to read its sequence. Nat. Struct. Mol. Biol. 29, 395–402 (2022).

22. Shipman, S. L., Nivala, J., Macklis, J. D. & Church, G. M. Molecular recordings by directed CRISPR spacer acquisition. Science 353, aaf1175 (2016).

23. Ramachandran, A., Summerville, L., Learn, B. A., DeBell, L. & Bailey, S. Processing and integration of functionally oriented prespacers in the Escherichia coli CRISPR system depends on bacterial host exonucleases. J. Biol. Chem. 295, 3403–3414 (2020).

24. Shiimori, M., Garrett, S. C., Graveley, B. R. & Terns, M. P. Cas4 nucleases define the PAM, length, and orientation of DNA fragments integrated at CRISPR loci. Mol. Cell 70, 814–824.e6 (2018).

25. Makarova, K. S. et al. Evolutionary classification of CRISPR–Cas systems: a burst of class 2 and derived variants. Nat. Rev. Microbiol. 18, 67–83 (2019).

26. Kieper, S. N. et al. Cas4 facilitates PAM-compatible spacer selection during CRISPR adaptation. Cell Rep. 22, 3377–3384 (2018).

27. Lee, H., Zhou, Y., Taylor, D. W. & Sashital, D. G. Cas4-dependent prespacer processing ensures high-fidelity programming of CRISPR arrays. Mol. Cell 70, 48–59.e5 (2018).

28. Kim, S. et al. Selective loading and processing of prespacers for precise CRISPR adaptation. Nature 579, 141–145 (2020).

29. Drabavicius, G. et al. DnaQ exonuclease-like domain of Cas2 promotes spacer integration in a type I-E CRISPR-Cas system. EMBO Rep. 19, (2018).

30. Lee, H., Dhingra, Y. & Sashital, D. G. The Cas4-Cas1-Cas2 complex mediates precise prespacer processing during CRISPR adaptation. Elife 8, (2019).

31. Hu, C. et al. Mechanism for Cas4-assisted directional spacer acquisition in CRISPR–Cas. Nature 598, 515–520 (2021).

32. Hudaiberdiev, S. et al. Phylogenomics of Cas4 family nucleases. BMC Evol. Biol. 17, 232 (2017).

33. Dhingra, Y., Suresh, S. K., Juneja, P. & Sashital, D. G. PAM binding ensures orientational integration during Cas4-Cas1-Cas2-mediated CRISPR adaptation. Mol. Cell 82, 4353–4367.e6 (2022).

34. Shiriaeva, A. A. et al. Host nucleases generate prespacers for primed adaptation in the E. coli type I-E CRISPR-Cas system. Sci Adv 8, eabn8650 (2022).

35. Makarova, K. S. & Koonin, E. V. Annotation and classification of CRISPR-Cas systems. Methods in Molecular Biology 47–75 (2015).

36. Nuñez, J. K. et al. Cas1–Cas2 complex formation mediates spacer acquisition during CRISPR–Cas adaptive immunity. Nat. Struct. Mol. Biol. 21, 528–534 (2014).

37. Wright, A. V. et al. Structures of the CRISPR genome integration complex. Science 357, 1113–1118 (2017).

38. Punjani, A. & Fleet, D. J. 3D variability analysis: Resolving continuous flexibility and discrete heterogeneity from single particle cryo-EM. J. Struct. Biol. 213, 107702 (2021).

39. Jakhanwal, S. et al. A CRISPR-Cas9–integrase complex generates precise DNA fragments for genome integration. Nucleic Acids Res. 49, 3546–3556 (2021).

40. Wright, A. V., Nuñez, J. K. & Doudna, J. A. Biology and applications of CRISPR systems: harnessing nature’s toolbox for genome engineering. Cell 164, 29–44 (2016).

41. Nuñez, J. K., Bai, L., Harrington, L. B., Hinder, T. L. & Doudna, J. A. CRISPR immunological memory requires a host factor for specificity. Mol. Cell 62, 824–833 (2016).

42. Budhathoki, J. B. et al. Real-time observation of CRISPR spacer acquisition by Cas1-Cas2 integrase. Nat. Struct. Mol. Biol. 27, 489–499 (2020).

43. Shipman, S. L., Nivala, J., Macklis, J. D. & Church, G. M. CRISPR–Cas encoding of a digital movie into the genomes of a population of living bacteria. Nature 547, 345–349 (2017).

44. Bhattarai-Kline, S. et al. Recording gene expression order in DNA by CRISPR addition of retron barcodes. Nature 608, 217–225 (2022).

## Methods References

45. Punjani, A., Rubinstein, J. L., Fleet, D. J. & Brubaker, M. A. cryoSPARC: algorithms for rapid unsupervised cryo-EM structure determination. Nat. Methods 14, 290–296 (2017).

46. Bepler, T. et al. Positive-unlabeled convolutional neural networks for particle picking in cryo-electron micrographs. Nat. Methods 16, 1153–1160 (2019).

47. Punjani, A., Zhang, H. & Fleet, D. J. Non-uniform refinement: adaptive regularization improves single-particle cryo-EM reconstruction. Nat. Methods 17, 1214–1221 (2020).

48. Jumper, J. et al. Highly accurate protein structure prediction with AlphaFold. Nature 596, 583–589 (2021).

49. Pettersen, E. F. et al. UCSF ChimeraX: Structure visualization for researchers, educators, and developers. Protein Sci. 30, 70–82 (2021).

50. Emsley, P., Lohkamp, B., Scott, W. G. & Cowtan, K. Features and development of Coot. Acta Crystallogr. D Biol. Crystallogr. 66, 486–501 (2010).

51. Liebschner, D. et al. Macromolecular structure determination using X-rays, neutrons and electrons: recent developments in Phenix. Acta Crystallogr D Struct Biol 75, 861–877 (2019).

52. Biswas, A., Staals, R. H. J., Morales, S. E., Fineran, P. C. & Brown, C. M. CRISPRDetect: A flexible algorithm to define CRISPR arrays. BMC Genomics 17, 356 (2016).

53. Steinegger, M. et al. HH-suite3 for fast remote homology detection and deep protein annotation. BMC Bioinformatics 20, 473 (2019).

54. Meng, E. C., Pettersen, E. F., Couch, G. S., Huang, C. C. & Ferrin, T. E. Tools for integrated sequence-structure analysis with UCSF Chimera. BMC Bioinformatics 7, 339 (2006).

