## Supplemental Tables 1-3, Extended Data Figures 1-10 for "Genome expansion by a CRISPR trimmer-integrase"

**Supplemental Movie 1**. **Visualization of repeat DNA flexibility of the Cas1:Cas2-DEDDh half-integration complex using 3-D variability analysis.** The PAM side is shown. See Materials and Methods for further detail.

**Supplementary Table 1. Sequences of the type I-E *Megasphaera NM10_related* Cas1 and Cas2-DEDDh proteins used in this study.**

| Protein | Sequence |
|---|---|
| Cas1 | MAGPIIAGKSESSELPRVEDRATFIYIEHAKINRVDSAVTVAEAKG<br>VVRIPAAMIGVLLLGPGTDISHRAVELLGDTGTALVWVGEQGVR<br>YYASGRALARSTRFLVKQAELVTNERSRLRVARRMYQMRFPTE<br>DVSKLTMQQLRSHEGARVRRKYRELSKKYNVPWKKRVYNPDD<br>FAGGDPINQALSAAHVALYGLVHSVVAALGLSPGLGFVHTGHDR<br>SFIYDVADLYKAEITVPIAFAVAAEAEEGQDIGQLARLRTRDAFV<br>DGKILKRMVKDLQTLLEIPEEGQIEAEPLSLWDDKEKLVPYGVN<br>YSEVTSCP* |
| Cas2-<br>DEDDh | MPMTVITLKNVPQSLRGDLTRWMQEIATGVYVGNFNSRIREYLW<br>RRVQETMGAGEASMCFAARNELGYDFLTENASRSVIDYDGLPLI<br>FIPKEQSAVSDLPKGFSTAAKLHRAHIAGSGKKKEKPIRYVVIDIE<br>TDGKDAKRNHILEIGAIRCEDGKETHFTALISGDAVPPSITKLTGIT<br>ATLLQKEGQEEKKVLTAFREFIGDDDLVGYHVSFDIEFLRQAFKK<br>YGLGYLKNKTHDLLRIVKKEQLFQADYKLETSLQSYGIHKKVPH<br>RALGDAELVKCLAKKLNKF* |

**Supplementary Table 2. DNA oligonucleotides used in this study.** * indicates phosphorothioate bond.

| Name/Description | Sequence | Figure |
|---|---|---|
| S1_top | actcatgatgcacaagtggttgcgcgtg | 1 |
| S1_bottom | gcaaccacttgtgcatcatgagtgatga | 1 |
| S2_top<br><br>mixed OH 1 (AA PAM)_top | actcatgatgcacaagtggttgcgcgtgAACCCAGTTG | 1, ED1, ED2, ED3 |
| S2_bottom<br><br>Bottom strand of prespacers<br><br>15-nt 3’OH / 23-bp duplex / 15-nt 3’OH (bottom) | gcaaccacttgtgcatcatgagtgatgaAACCCAGTTG | 1, 4, 5, ED1, ED2, ED3, ED10 |
| S3_top | actcatgatgcacaagtggttgcgcgtgACCTAGAAGTAACCCAGTTG | ED2 |
| S3_bottom | gcaaccacttgtgcatcatgagtgatgaACCTAGAAGTAACCCAGTTG | ED2 |
| polyA OH (AA PAM)_top | actcatgatgcacaagtggttgcgcgtgAAAAAAAAAA | 1, ED3 |
| poly T OH (TT PAM)_top | actcatgatgcacaagtggttgcgcgtgTTTTTTTTTT | 1, ED3 |
| poly C OH (CC PAM)_top | actcatgatgcacaagtggttgcgcgtgCCCCCCCCCC | 1, ED3 |
| poly G OH (GG PAM)_top | actcatgatgcacaagtggttgcgcgtgGGGGGGGGGG | 1, ED3 |
| mixed OH 2 (TT PAM)_top | actcatgatgcacaagtggttgcgcgtgTTTTCAGTTG | 1, ED3 |
| mixed OH 3 (TT PAM)_top<br><br>15-nt 3’OH / 23-bp duplex / 15-nt 3’OH (top) | actcatgatgcacaagtggttgcgcgtgTTCCCAGTTG | 1, ED3, ED10 |
| mixed OH 4 (TA PAM)_top | actcatgatgcacaagtggttgcgcgtgTACCCAGTTG | 1, ED3 |
| Full-site prespacer_top<br><br>Full-site prespacer | aaacggagacctggtctcaatctgcgtgTTCCCAGTTG | 4, 5, ED9 |
| Full-site prespacer_bottom<br><br>Full-site prespacer (atg barcode)_bottom | agattgagaccaggtctccgtttcatgaAACCCAGTTG | 4, 5, ED9 |
| CryoEM prespacer PAM-deficient strand | actcatgatgcacaagtggttgcgcgtg*AACCCAGTTG | 2 |
| CryoEM prespacer PAM-deficient complement strand | gcaaccacttgtgcatcatgagtgatga*AACCCAGTTG | 2 |

| | | |
|---|---|---|
| CryoEM prespacer PAM-containing strand | actcatgatgcacaagtggttgcgcgtgTTCCC*AGTTG | 2 |
| CryoEM prespacer PAM-containing strand complement | gcaaccacttgtgcatcatgagtgatga*AACCCAGTTG | 2 |
| Half-site substrate bottom strand (spacer-repeat-leader) | tgcgcgtgggatcacccccgctcgtgcgggaaagacagtaatggattcctttattttcgcccttttacgcttactgac*g*t | 3, ED6 |
| Half-site substrate 1 leader strand | acgtcagtaagcgtaaaagggcgaaaataaaggaatccatta*c*t | 3, ED6 |
| Half-site substrate protospacer-repeat-spacer strand | agattgagaccaggtctccgtttcatgagtctttcccgcacgagcgggggtgatcccacgcg*c*a | 3, ED6 |
| Half-site substrate 1 protospacer + unprocessed PAM strand | aaacggagacctggtctcaatctgcgtgTTCCC | 3, ED6 |
| Half-site substrate 2 protospacer strand | aaacggagacctggtctcaatctgcgtg | 3, ED6 |
| CryoEM half-site substrate protospacer strand | aaacggagacctggtctcaatctgcgtgT*T*CCC | 3 |
| Full-site prespacer_top (with PAM)<br><br>Full-site prespacer (tcg barcode, PAM)_top | aaacggagacctggtctcaatctgtcggTTCCCAGTTG | 4, ED9 |
| Full-site prespacer_top (no PAM)<br><br>Full-site prespacer (tcg barcode, no PAM)_top | aaacggagacctggtctcaatctgtcggAACCCAGTTG | 4, ED9 |
| Full-site prespacer_bottom<br><br>Full-site prespacer (gat barcode)_bottom | agattgagaccaggtctccgtttcgataAACCCAGTTG | 4, ED9 |
| Full-site prespacer (cgt barcode, no PAM)_top | aaacggagacctggtctcaatctgcgtgAACCCAGTTG | 4, ED9 |
| D1_top<br><br>SS1 | CTGCATCTGGGTATCATCACTCATGATGCACTAGTGGATGCGCGTGATCCTATGCATGA | 5 |
| D1_bottom | TCATGCATAGGATCACGCGCATCCACTAGTGCATCATGAGTGATGATACCCAGATGCAG | 5 |
| D2_top<br><br>SS2 | CTGCATCTGGGTATCATCACTCATGATGCACTAGTGGATGCGCGTGTTCCTATGCATGA | 5 |
| D2_bottom | TCATGCATAGGAACACGCGCATCCACTAGTGCATCATGAGTGATGATACCCAGATGCAG | 5 |
| SS3 | CTGCATCTGGGTATCATCACTCATGATGCACTAGTGGATGTTCGTGATCCTATGCATGA | 5 |
| SS4 | CTGCATCTGGGTATCATCACTCATGATGCACTA | 5 |

| | | |
|---|---|---|
| | GTTGATGCGCGTGATCCTATGCATGA | |
| 11-nt 3'OH / 31-bp duplex / 11-nt 3'OH (top) | catcactcatgatgcacaagtggttgcgcgtgTTCCCAGTTG | ED10 |
| 11-nt 3'OH / 31-bp duplex / 11-nt 3'OH (bottom) | acgcgcaaccacttgtgcatcatgagtgatgaAACCCAGTTG | ED10 |
| 13-nt 3'OH / 27-bp duplex / 13-nt 3'OH (top) | tcactcatgatgcacaagtggttgcgcgtgTTCCCAGTTG | ED10 |
| 13-nt 3'OH / 27-bp duplex / 13-nt 3'OH (bottom) | gcgcaaccacttgtgcatcatgagtgatgaAACCCAGTTG | ED10 |
| 17-nt 3'OH / 19-bp duplex / 17-nt 3'OH (top) | tcatgatgcacaagtggttgcgcgtgTTCCCAGTTG | ED10 |
| 17-nt 3'OH / 19-bp duplex / 17-nt 3'OH (bottom) | aaccacttgtgcatcatgagtgatgaAACCCAGTTG | ED10 |
| 19-nt 3'OH / 15-bp duplex / 19-nt 3'OH (top) | atgatgcacaagtggttgcgcgtgTTCCCAGTTG | ED10 |
| 19-nt 3'OH / 15-bp duplex / 19-nt 3'OH (bottom) | ccacttgtgcatcatgagtgatgaAACCCAGTTG | ED10 |
| Chloramphenicol selection cassette PCR forward primer | GGCCGGTCTCCAGATtgatcggcacgtaagaggttc | 4, ED10 |
| Chloramphenicol selection cassette PCR reverse primer | GGCCTGGTCTCAAAACattctcaccaataaaaaacgcccg | 4, ED10 |

#### Table S3: Cryo-EM Map and Model Validation

| Model | Model 1 (no PAM) | Model 2 (PAM) | Model 3 (PAM-DEDDh) | Model 4 (Half-site) | Model 5 (Halfi-site linear) | Model 6 (Half-site bent) |
|---|---|---|---|---|---|---|
| **Data collection and processing** | | | | | | |
| Electron Microscope | Talos Arctica | | | | | |
| Electron Detector | K3 Direct Electron Detector | | | | | |
| Magnification | x36,000 | | | | | |
| Voltage (kV) | 200 | | | | | |
| Electron dose (e-/Å$^{2}$) | 50 | | | | | |
| Pixel size (Å) | 1.115 | | | | | |
| Defocus range | 0.8 - 2.2 µm | 0.0 - 2.2 µm | | 0.8 - 2.2 µm | | |
| Tilt angle | 0° | 20° | | 20° | | |
| Grids | carbon 2/2 300 mesh C-flat grids | 1.2/1.3 300 mesh UltrAuFoil gold grids | | 1.2/1.3 300 mesh UltrAuFoil gold grids | | |
| Complex molarity | 8 µM | 2.4 µM | | 2.5 µM | | |
| **3D reconstruction** | | | | | | |
| Raw images | 2220 | 1457 | 1457 | 6066 | 6066 | 6066 |
| Initial particles | 701,623 | 3,101,776 | 228,220 | 1,836,610 | 1,048,353 | 1,836,610 |
| Final particles | 461,266 | 1,420,721 | 49,383 | 1,048,353 | 58,475 | 53,545 |
| Map resolution (Å) | 3.13 | 2.91 | 3.53 | 3.14 | 4.06 | 3.88 |
| FSC threshold | 0.143 | 0.143 | 0.143 | 0.143 | 0.143 | 0.143 |
| Map sharpening B-factor (Å$^{-2}$) | 140 | 138 | 112.1 | 165.7 | 118.7 | 142.6 |
| **Model refinement** | | | | | | |
| Initial model used | Model 2 | *Ab initio AlphaFold2 model* | Model 2 | Model 2 | Model 2 | Model 2 |
| Model resolution | 3.3 | 3.1 | 4.1 | 3.3 | 4.4 | 4.3 |
| FSC threshold | 0.5 | 0.5 | 0.5 | 0.5 | 0.5 | 0.5 |
| <u>Model composition</u> | | | | | | |
| Nonhydrogen atoms | 10742 | 11433 | 12724 | 13486 | 14567 | 13612 |
| Protein residues | 1228 | 1304 | 1461 | 1359 | 1474 | 1325 |
| Nucleotide | 56 | 59 | 61 | 136 | 145 | 156 |
| Ligands | 0 | 0 | 0 | 0 | 0 | 0 |
| <u>B factors (Å$^{2}$)</u> | | | | | | |
| Protein | 65.37 | 55.33 | 154.31 | 61.24 | 155.43 | 127.03 |
| Nucleotide | 76.58 | 48.17 | 142.08 | 132.98 | 200.4 | 227.2 |
| <u>R.m.s. deviations</u> | | | | | | |
| Bond length (Å) | 0.003 | 0.003 | 0.003 | 0.004 | 0.003 | 0.003 |
| Bond angles (°) | 0.521 | 0.511 | 0.647 | 0.546 | 0.585 | 0.547 |
| **Validation** | | | | | | |
| MolProbity score | 1.71 | 1.73 | 1.85 | 1.78 | 1.92 | 1.95 |
| Clash score | 10.69 | 12.06 | 15.87 | 10.52 | 17.83 | 14.49 |
| Rotamer outlier (%) | 0 | 0.09 | 0.08 | 0 | 0.08 | 0 |
| <u>Ramachandran statistics</u> | | | | | | |
| Favored | 97.11 | 97.28 | 97.22 | 96.44 | 97.05 | 95.88 |
| Allowed | 2.89 | 2.72 | 2.78 | 3.56 | 2.88 | 4.12 |
| Outlier | 0 | 0 | 0 | 0 | 0.07 | 0 |
| Rama-Z score, whole (r.m.s. Rama-Z) | 1.05 (0.24) | 1.24 (0.24) | 0.14 (0.22) | 0.77 (0.23) | 0.72 (0.21) | -0.05 (0.23) |
| Map CC (box) | 0.7 | 0.68 | 0.69 | 0.69 | 0.82 | 0.81 |
| Map CC (mask) | 0.77 | 0.78 | 0.72 | 0.78 | 0.74 | 0.79 |

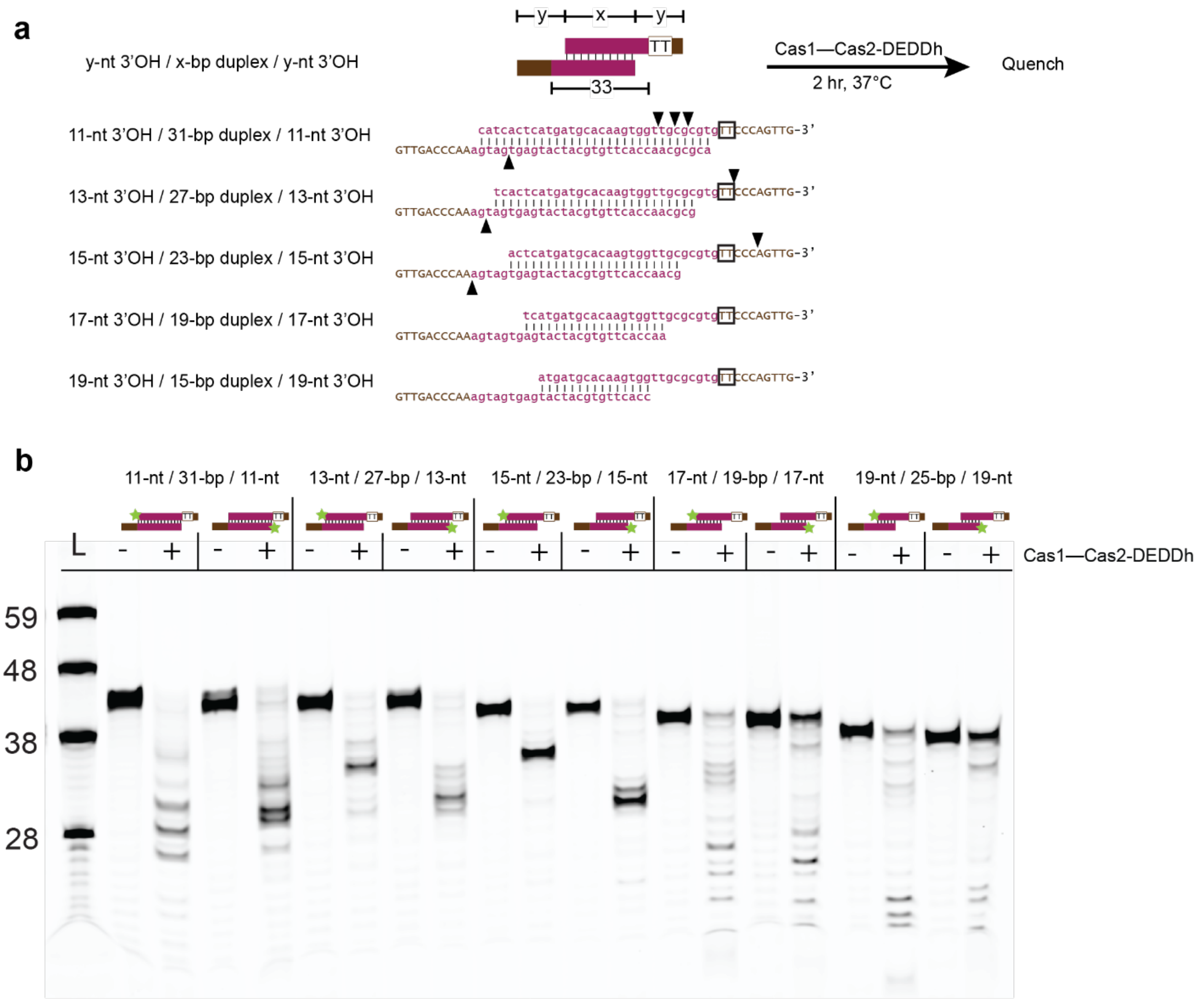


**Extended Data Figure 1. Cas1—Cas2-DEDDh exhibits strict requirement for pre-duplexed 23-bp DNA duplex prespacer for functional ruler-guided trimming. (a)** Prespacer substrates with varying duplex lengths used in processing assay. The duplex and overhang lengths are indicated, and the TT PAM motif is boxed. The arrowheads indicate specific processing positions as observed from the processing products visualized on the denaturing gel in **b**. **(b)** Processing assay with pre-duplexed prespacer substrates shown in **a**. Substrates are schematized.

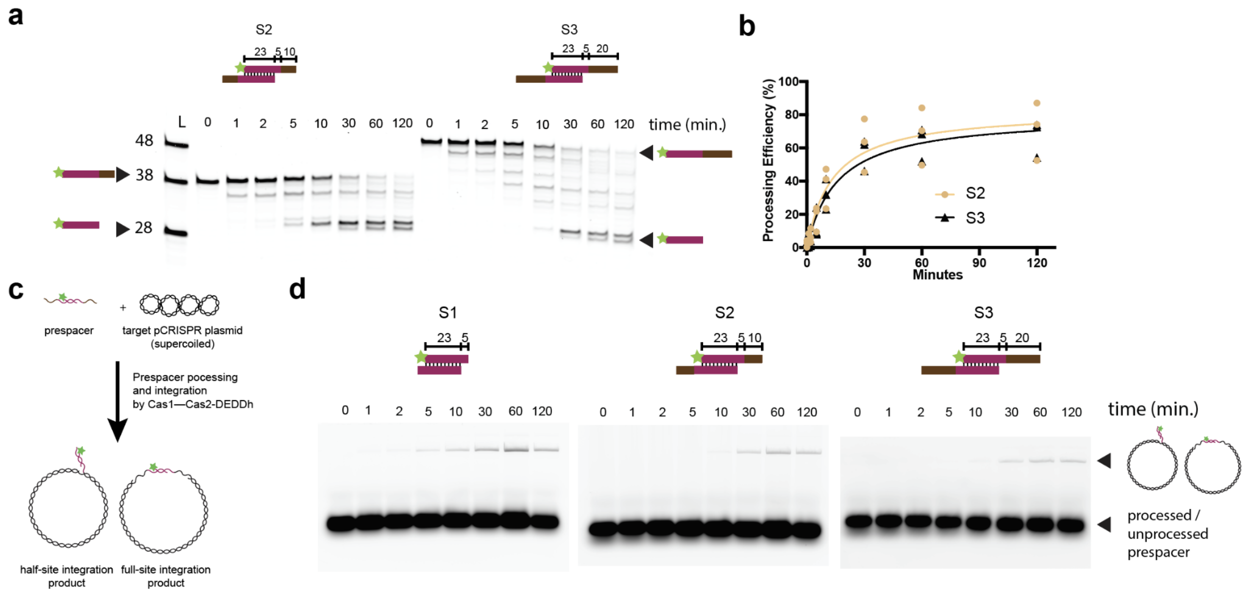


**Extended Data Figure 2. Kinetics of exonuclease trimming and ligation by Cas1:Cas2-DEDDh *in vitro*. (a)** Time-course reactions of ruler-guided trimming by Cas1:Cas2-DEDDh using substrates S2 and S3 over two hours. **(b)** Quantification of time-course reactions of ruler-guided trimming shown in **a**. Processing efficiency is calculated as the percentage of fully processed product formation at 28-29 nt (n = 3 biologically independent experiments). **(c)** Schematic of *in vitro* ligation assay with integration target pCRISPR plasmid and prespacer substrate. Star indicates 6-carboxyfluorescein (6-FAM) label. **(d)** Time-course reactions of ligation assay with substrates from (b) over 2 hours. Note formation of a 6-FAM-labeled pCRISPR plasmid ligation product band.

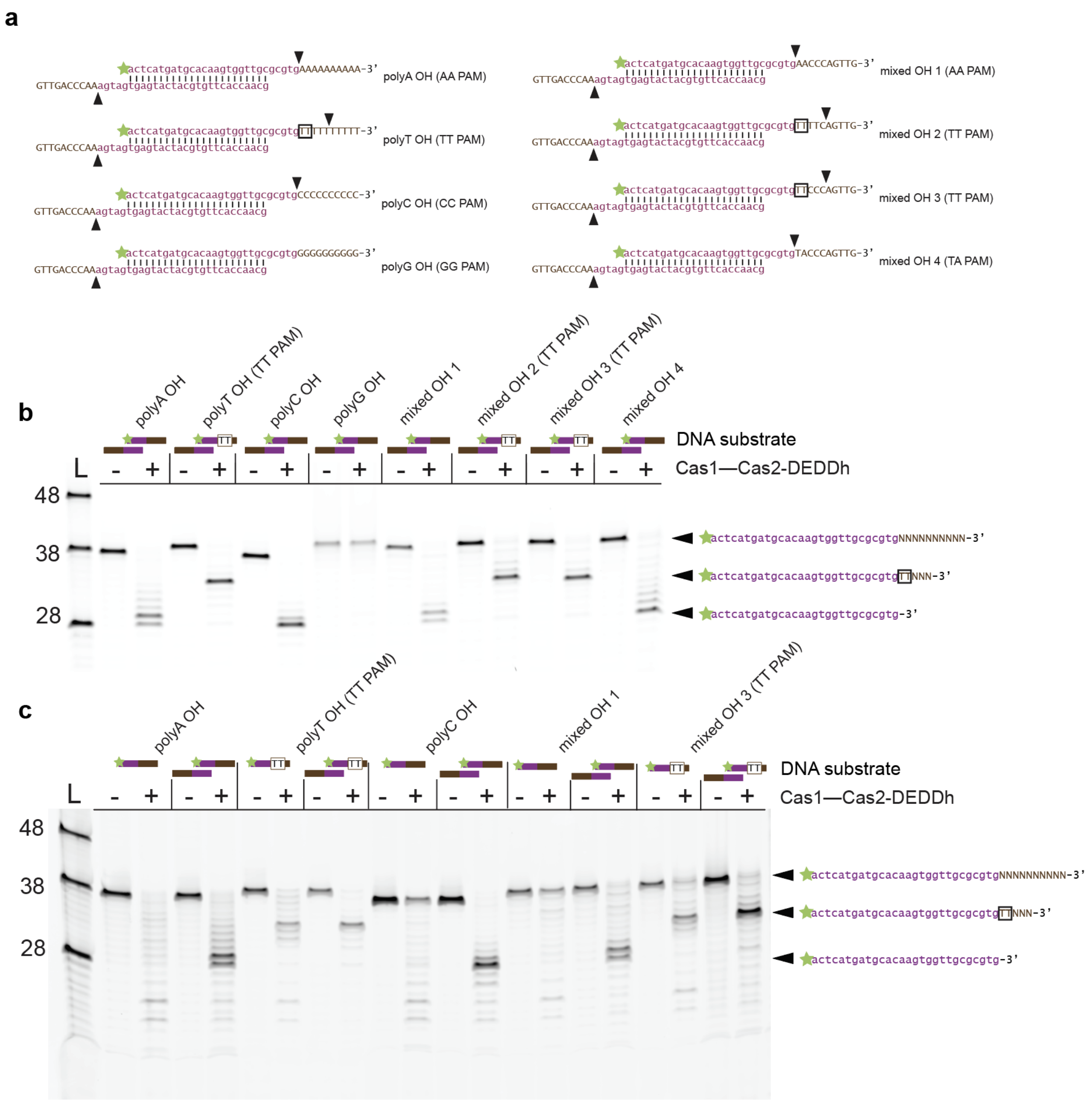


**Extended Data Figure 3. Complete PAM-mediated trimming experiments. (a)** Substrates used in processing assay with varied 3' overhang sequences. TT PAM is boxed. Arrowhead indicates how far substrate strand gets trimmed during processing reaction. Star indicates 6-carboxyfluorescein label. **(b)** Processing assay with substrates shown in **a**. Substrates are schematized and TT PAM is indicated. Processing products are depicted on the right, TT PAM is boxed. **(c)** Processing assay with select substrates in **a** and their single-stranded DNA controls, TT PAM is boxed.

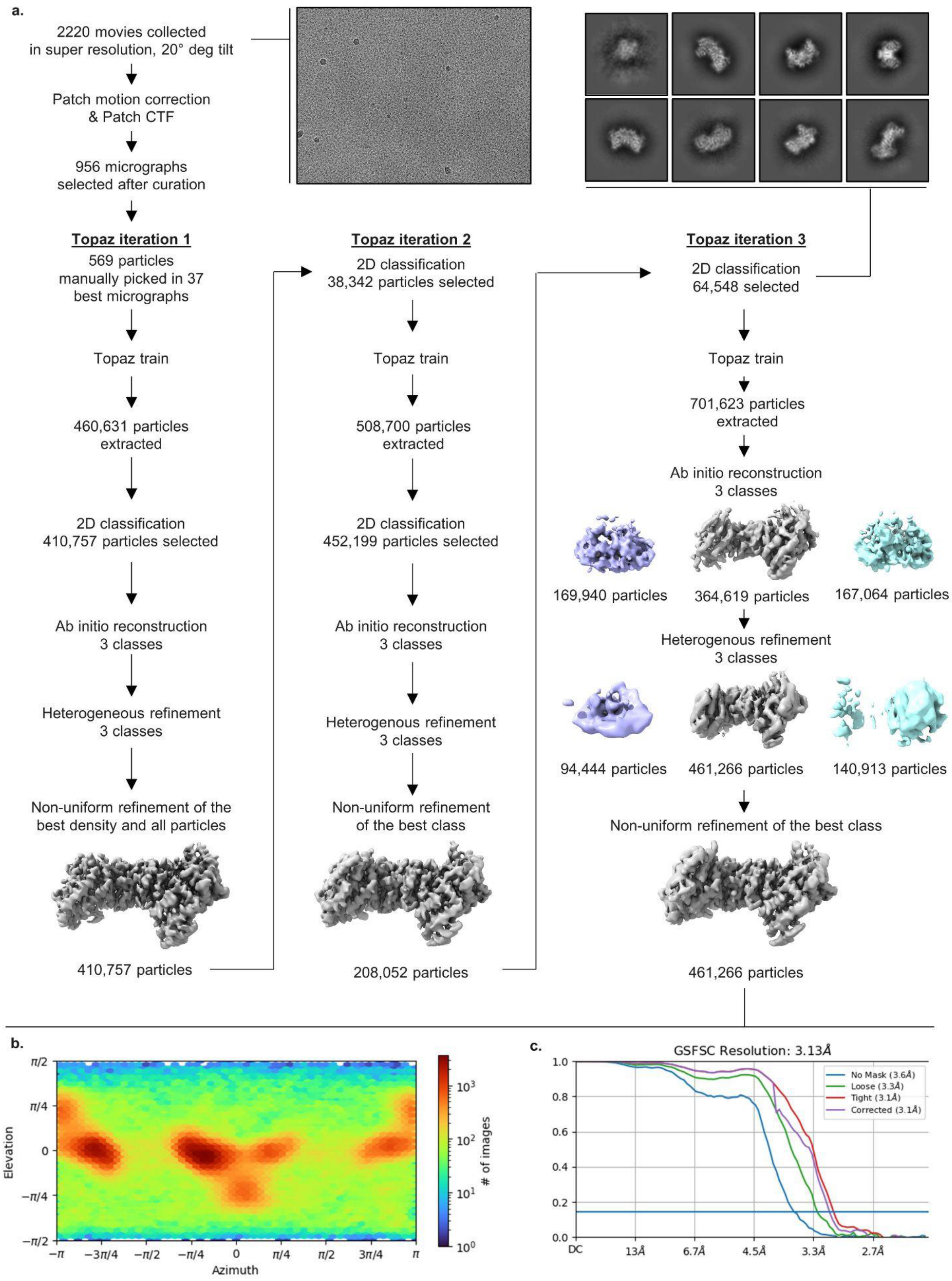
a.
2220 movies collected in super resolution, 20° deg tilt
Patch motion correction & Patch CTF
956 micrographs selected after curation
Topaz iteration 1
569 particles manually picked in 37 best micrographs
Topaz train
460,631 particles extracted
2D classification 410,757 particles selected
Ab initio reconstruction 3 classes
Heterogeneous refinement 3 classes
Non-uniform refinement of the best density and all particles
410,757 particles
Topaz iteration 2
2D classification 38,342 particles selected
Topaz train
508,700 particles extracted
2D classification 452,199 particles selected
Ab initio reconstruction 3 classes
Heterogenous refinement 3 classes
Non-uniform refinement of the best class
208,052 particles
Topaz iteration 3
2D classification 64,548 selected
Topaz train
701,623 particles extracted
Ab initio reconstruction 3 classes
169,940 particles
364,619 particles
167,064 particles
Heterogenous refinement 3 classes
94,444 particles
461,266 particles
140,913 particles
Non-uniform refinement of the best class
461,266 particles
b.
Elevation
Azimuth
### of images
c.
GSFSC Resolution: 3.13Å
No Mask (3.6Å)
Loose (3.3Å)
Tight (3.1Å)
Corrected (3.1Å)
DC
13Å
6.7Å
4.5Å
3.3Å
2.7Å

35

**Extended Data Figure 4. Flow-chart of the cryo-EM single particle reconstruction of the PAM-deficient prespacer bound Cas1-Cas2/DEDDh. (a)** Map generation pipeline in cryoSPARC consisting of three Topaz training[46] iterations, including representative 2D class averages and 3D maps resulting from *ab initio* reconstruction and further heterogeneous, non-uniform refinement. **(b)** Orientation distribution of the final set of refined particles. **(c)** Gold standard FSC curve of the atomic model refined to the final cryoSPARC sharp map.

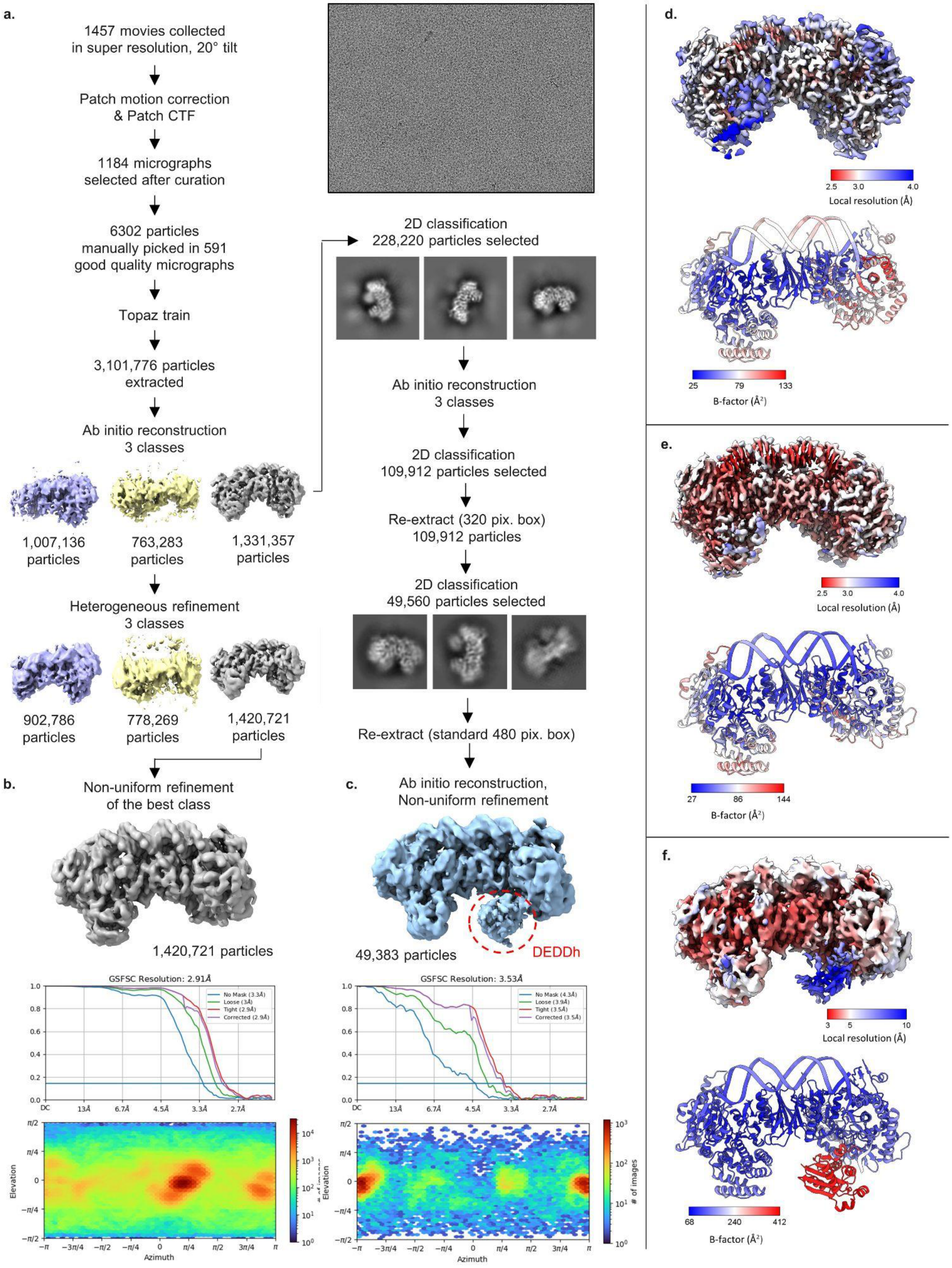
a.
1457 movies collected
in super resolution, 20° tilt
Patch motion correction
& Patch CTF
1184 micrographs
selected after curation
6302 particles
manually picked in 591
good quality micrographs
Topaz train
3,101,776 particles
extracted
Ab initio reconstruction
3 classes
1,007,136
particles
763,283
particles
1,331,357
particles
Heterogeneous refinement
3 classes
902,786
particles
778,269
particles
1,420,721
particles
2D classification
228,220 particles selected
Ab initio reconstruction
3 classes
2D classification
109,912 particles selected
Re-extract (320 pix. box)
109,912 particles
2D classification
49,560 particles selected
Re-extract (standard 480 pix. box)
b.
Non-uniform refinement
of the best class
1,420,721 particles
GSFSC Resolution: 2.91Å
No Mask (3.3Å)
Loose (3Å)
Tight (2.9Å)
Corrected (2.9Å)
Elevation
Azimuth
c.
Ab initio reconstruction,
Non-uniform refinement
49,383 particles
DEDDh
GSFSC Resolution: 3.53Å
No Mask (4.3Å)
Loose (3.9Å)
Tight (3.5Å)
Corrected (3.5Å)
### of images
d.
Local resolution (Å)
B-factor (Å²)
e.
Local resolution (Å)
B-factor (Å²)
f.
Local resolution (Å)
B-factor (Å²)

**Extended Data Figure 5. Flow-chart of the cryo-EM single particle reconstruction of the PAM-containing prespacer bound Cas1-Cas2 and resolution of the DEDDh density. (a)** Cryo-EM data collection parameters and map generation pipeline in cryoSPARC, including representative 2D class averages and 3D maps resulting from *ab initio* reconstruction and further heterogeneous refinement. **(b)** 3D maps, orientation distribution, and gold standard FSC curve of the final cryoSPARC map for the PAM-only (**b**) and DEDDh-containing (**c**) densities. **(d-f)** Final sharp maps of Cas1:Cas2-DEDDh bound to prespacers colored according to local resolution as calculated by cryoSPARC, and the final refined models colored with B-factors as calculated by Phenix, for: (**d**) PAM-deficient density, (**e**) PAM-containing DEDDh-absent density, and (**f**) PAM-containing, DEDDh-containing density.

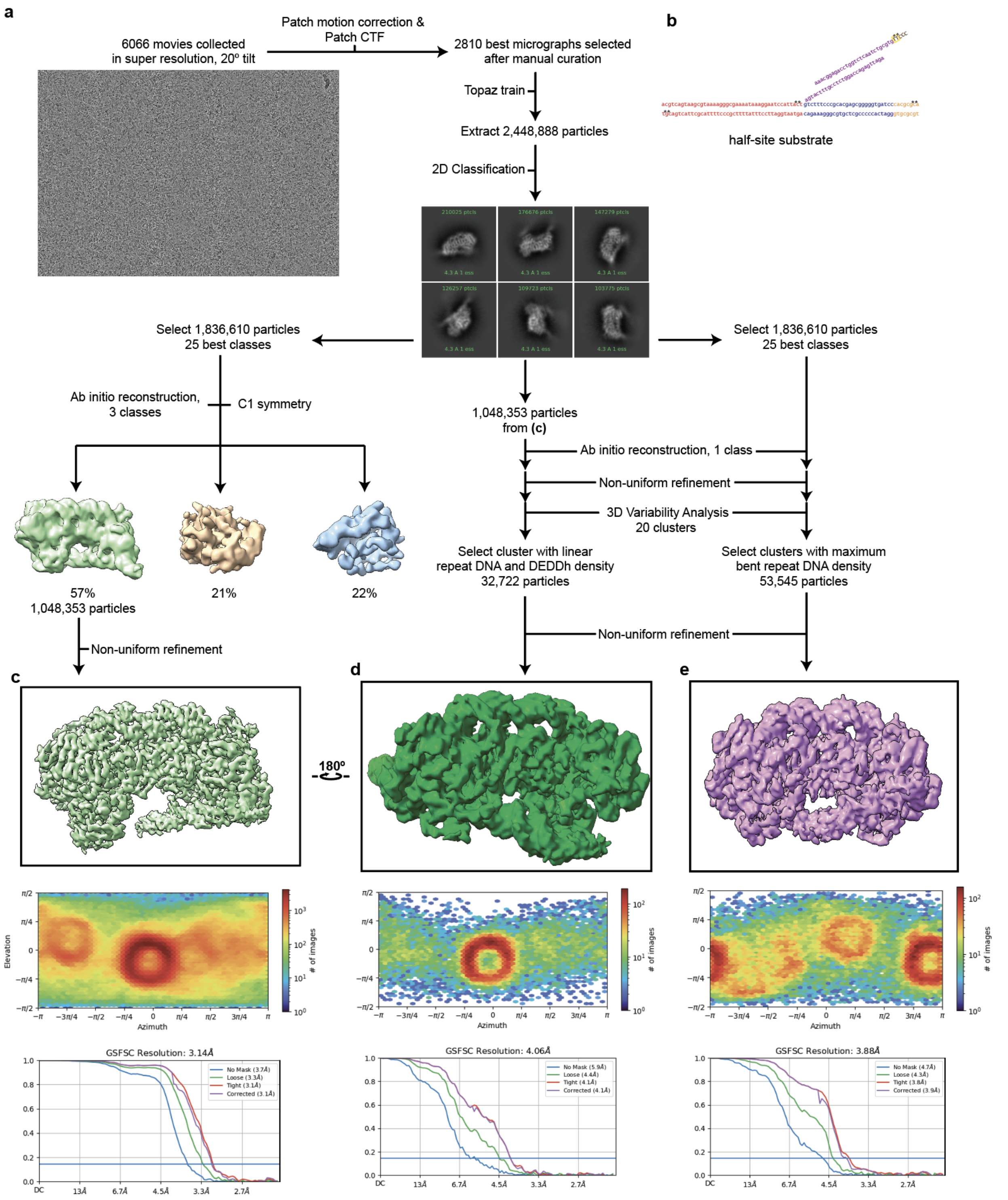


**Extended Data Figure 6. Flow-chart of the cryo-EM single particle reconstruction of the PAM-containing half-site substrate bound to Cas1-Cas2/DEDDh. (a)** Cryo-EM data collection parameters and map generation pipeline in cryoSPARC, including representative 2D class averages and 3D maps resulting from *ab initio* reconstruction and further heterogeneous refinement. Different particle stacks were used for generation of the high resolution structure **(c),** and 3D variability analysis resulting in linear **(d)** and bent (**e)** density maps. **(b)** Representation of the 4 oligonucleotide half integration construct used in this study. Asterisk indicates a phosphorothioate bond. **(c-e)** 3D maps, orientation distribution, and gold standard FSC curve of the atomic model refined to the final cryoSPARC sharp map for each half-site structure.

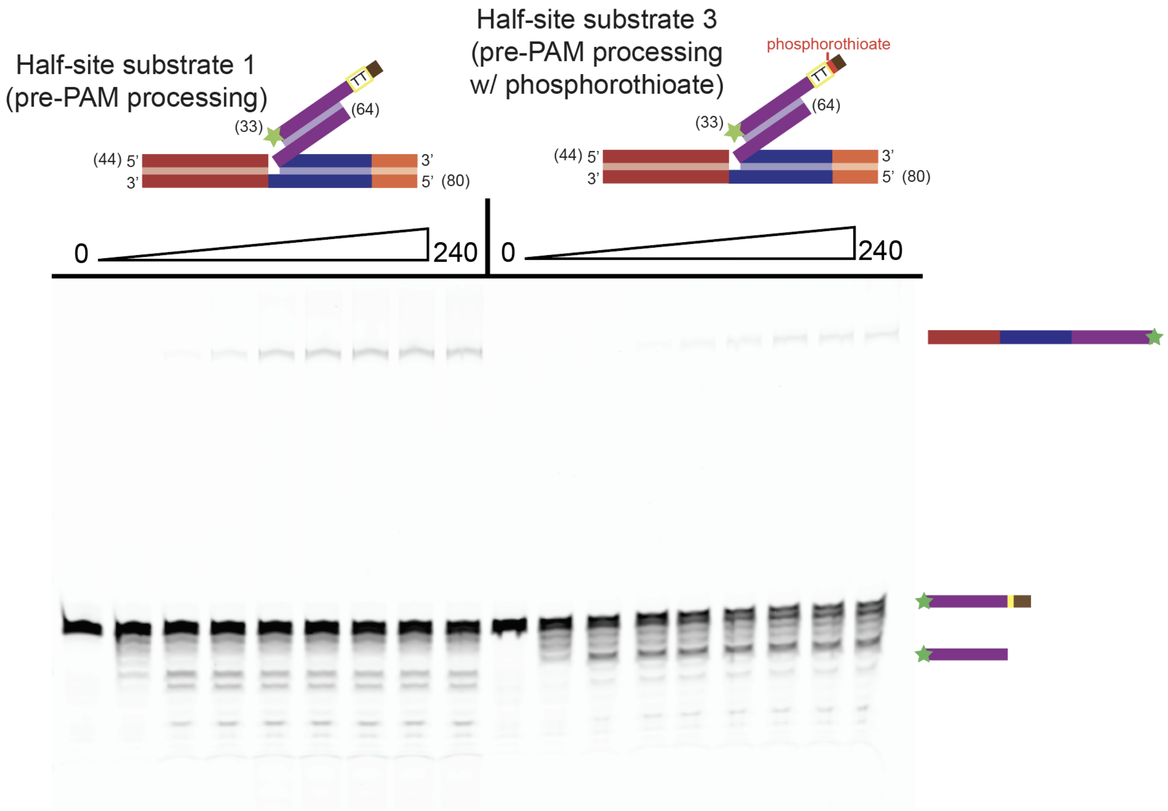


**Extended Data Figure 7. Incomplete resistance of phosphorothioate PAM nucleotides to DEDDh.** Processing assay (4 hour) with substrates shown in above. Substrates are schematized and TT PAM is indicated, along with phosphorothioation. Processing products are depicted on the right.

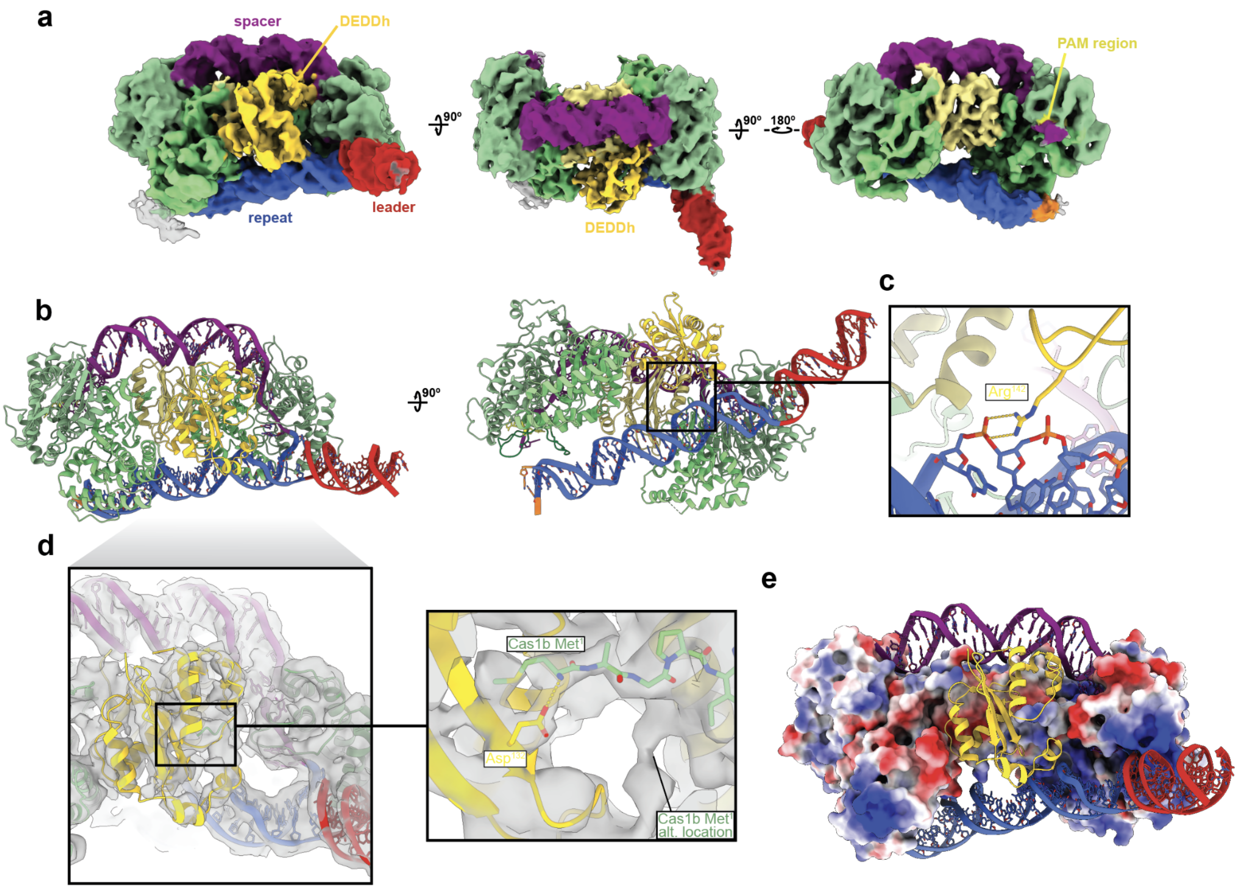


**Extended Data Figure 8. Molecular detail of Cas1:Cas2-DEDDh in the linear CRISPR repeat DNA conformation.** (**a**) Orthogonal views of the final sharpened cryo-EM densities for Cas1:Cas2-DEDDh bound to half-site DNA intermediates containing a phosphorothioated TT PAM, colored to demonstrate domain locations. **(b)** Non-PAM side and bottom views of Cas1:Cas2-DEDDh bound to half-site intermediate DNA. **(c)** Detail depicting potential interaction between Arg$^{142}$ of DEDDh and CRISPR repeat DNA phosphate backbone. **(d)** Non-PAM side view of the DEDDh domain with the sharp map superimposed (threshold: 0.05). Right, detail in the catalytic pocket of DEDDh, with Asp$^{132}$ shown in close proximity to an extension attributed to the N-terminus of Cas1b. **(e)** Cas1:Cas2-DEDDh linear structure, with protein surfaces except the DEDDh domain colored according to electrostatic potential.

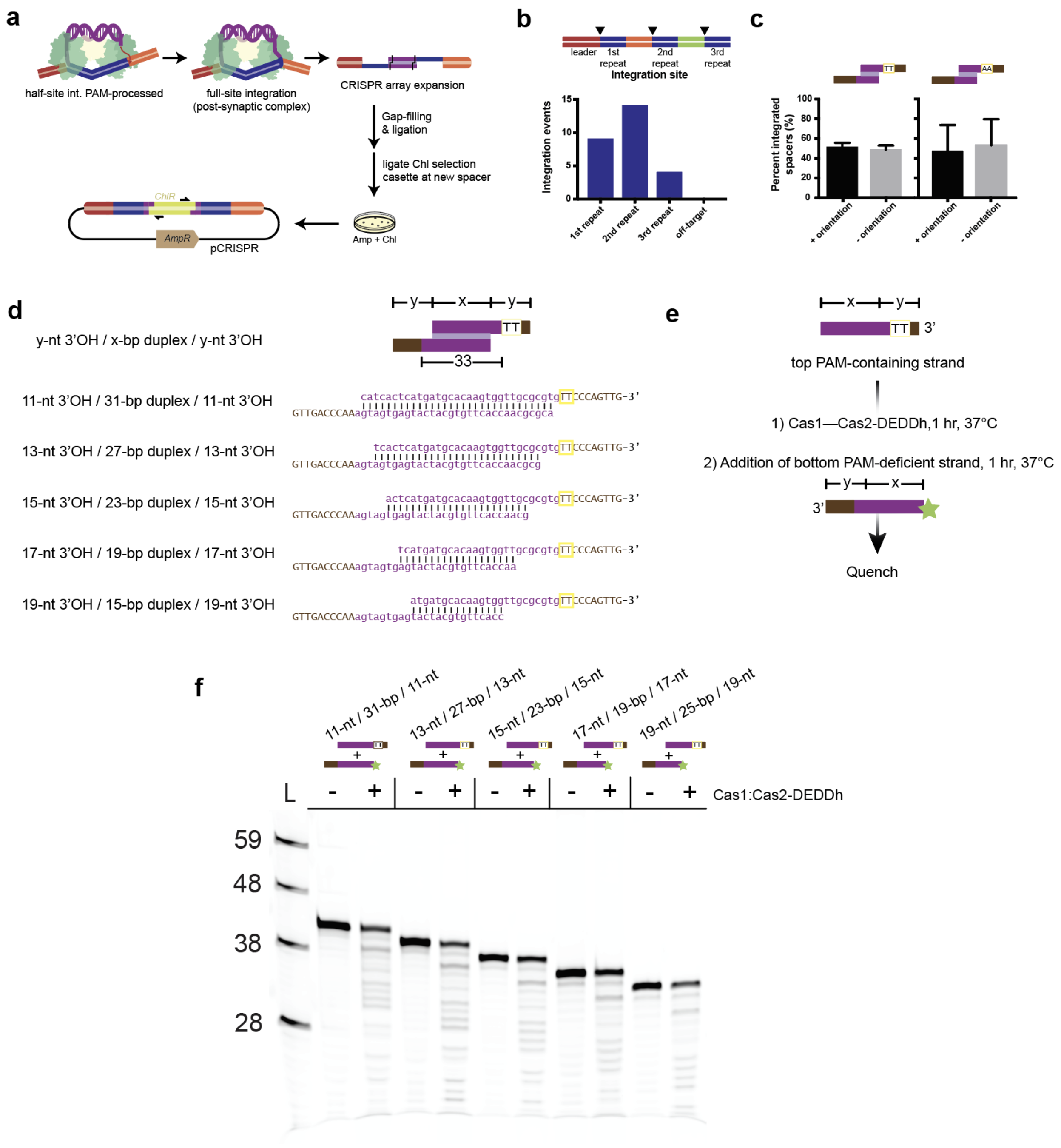


**Extended Data Figure 9. Integration site preference, orientation bias, and duplex requirements of Cas1—Cas2-DEDDh. (a)** Schematic depicting experimental workflow for full-site integration reconstitution. **(b)** Number of integration events at each integration site from sequenced clones. Integration sites are depicted by arrows at repeat borders. **(c)** Left, orientation of spacer insertion from prespacer containing PAM motif, + orientation orients original TT PAM containing end toward the leader, - orientation orients original TT PAM containing end away from the leader. Right, Orientation of spacer insertion from control prespacer without PAM motif (the TT PAM from left is replaced with AA). The mean and standard deviation of three independent replicates are shown. **(d)** Prespacer substrates with varying duplex lengths used in processing assay. The duplex and overhang lengths are indicated, and the TT PAM motif is boxed. **(e)** Schematic of processing assay with stepwise addition of the top and bottom prespacer strands of the substrates shown in **d**. The unlabeled top PAM-containing strand is incubated first with Cas1:Cas2-DEDDh followed by delayed addition of the labeled bottom PAM-deficient strand. **(f)** Processing assay with stepwise addition of the top and bottom prespacer strands of the substrates shown in **a.**.

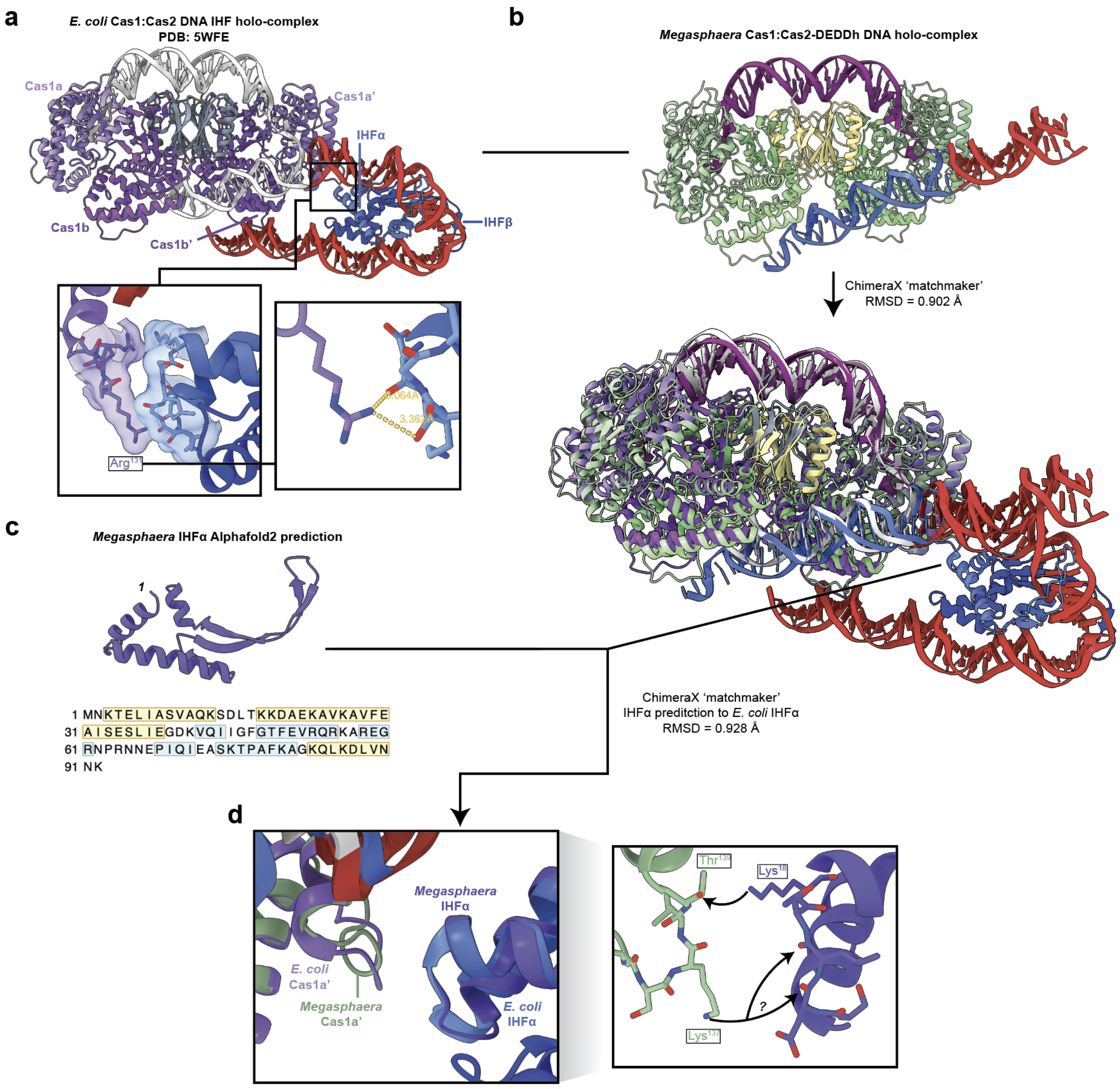


**Extended Data Figure 10. Comparison of Cas1:Cas2-DEDDh bound to the half-site intermediate with the complete *E. coli* IHF-containing integration complex. (a)** *E. coli* Cas1:Cas2 DNA IHF holo-complex (PDB: 5WFE). Subunit identities are indicated above. Inset boxes highlight the interface between Cas1a' and IHF. **(b)** Representation of the pipeline used to compare *Megasphaera* and *E. coli* integrases. The experimental structure solved in this work was superimposed with the *E. coli* complex using ChimeraX matchmaker[54]. **(c)** The sequence of a *Megasphaera* IHFα ortholog was found by protein BLAST of *E. coli* IHFα sequence in the

*Megasphaera* sp. An286 genome (Taxon ID: 1965622). The AlphaFold 2 sequence of the highest confidence hit is shown along with the protein sequence, with secondary structures indicated. **(d)** The *Megasphaera* IHFα was superimposed with the *E. coli* IHFα to approximate the interface in a hypothetical *Megasphaera* complex. Inset shows the structures overlaid (left) and with only *Megasphaera* proteins shown (right). Right, potential interactions at the interface of *Megasphaera* Cas1 and IHFα.
